# Galactic arms, and devil’s toenails: a synthesis of coiled morphology, applied to gastropods

**DOI:** 10.1101/2025.06.24.659690

**Authors:** Ido Filin

## Abstract

Biological structures lie at intersections of function, construction, and history. Conversely, the geometry of structures modulates how these three aspects of organic form manifest in different taxonomic groups and ecological circumstances. Here, I apply the familiar logarithmic conispiral model for coiled shells, to assess its fit to data on gastropods, and to study covariation in coiling behavior. Treating shell centerlines as *conical helices*, and drawing on analogies from astronomy and robotics, the expansion angle of the helix, and its helical slope, or lead angle, emerge as convenient parameters for studying shell growth and its optimization, trait covariation, and ontogenetic allometry. I demonstrate that by combining several published datasets. Recent arguments on the inadequacy of logarithmic helicospirals contain a known methodological pitfall. Adopting the conihelical parameterization circumvents the problem altogether, demonstrating good fits of conihelical centerlines to data. Further analysis of aperture inclinations, relative to conihelical centerlines, highlights the constructional developmental role of the environment (here, substrate) as a general principle of natural history, on par with adaptation. More broadly, ‘laws of form’ have practical use in data analyses and integration, and provide baseline references for ontogenetic allometry, growth gradients, and other morphological features (here, aperture inclination, and columellar folds).

## Introduction

Since antiquity, molluscan shells have served as exemplars for the immense variety of nature and, in the past two centuries, to how such diversity can be understood and unified by mathematical modeling (Moseley 1838, 1842; Thompson [1942] 1992; Raup 1961, 1966; Vermeij 1993; Stone 1996; McGhee 1999; Urdy 2015; Hammer 2016). Conchiferans also feature prominently in discussions of adaptation and constraint, the exchange among functional, constructional and historical aspects of form that is at the core of biology (Seilacher 1973; Savazzi 1990; Vermeij 1993, 2002; Vermeij & Sigwart 2026).

The standard point of departure for modeling coiled shelly organisms, both extant and fossil, is the logarithmic spiral, hereafter, *logspiral* (Thompson [1942] 1992; Raup 1961, 1966; McGhee 1999). Logspirals are famously self-similar, maintaining fixed proportions while expanding in size (isometric growth). Much literature discusses its inadequacy, due to commonly observed ontogenetic allometry (variation in coiling parameters with age or size) and irregular coiling, where even the coiling axis is not fixed, often associated with important life-history transitions (Cintra & de Souza Lopes 1952; Burnaby 1965, 1966; Davoli & Russo 1974; McGhee 1980, 1999, 2001; Okamoto 1988; Vermeij 1993; Savazzi 1990, 1996; Aldridge 1998; van Osselaer & Grosjean 2000; Liew *et al*. 2014; Chirat *et al*. 2021; Ohta *et al*. 2025).

Recently, Collins *et al*. (2021*b*) studied shell geometry from sectioned specimens and microtomography. Applying model selection, they concluded that logarithmic helicospirals are poor fit for gastropod shells. A first contribution of this study is to show that the conclusion of Collins *et al*. arises from the known ‘Origin of Longitudinal Coordinates’ methodological problem (van Osselaer & Grosjean 2000). When fully recognized and circumvented, logarithmic conispirals produce very good fits.

Beyond this reminder on methodological aspects of logspiral geometry, the “highly integrated morphology” (Monnet *et al*. 2009) of conispiral shells means that traits covary tightly, and thus translate to a low-dimensional morphospace. Consequently, there are many alternative equivalent ways to parameterize conispiral shells (Moseley 1838, 1842; Thompson [1942] 1992; Raup 1966; Løvtrup & von Sydow 1974; Ekaratne & Crisp 1983; Okamoto 1988; Ackerly 1989*a*; Rice 1998; Noshita 2014; Filin 2026*b*). Particularly, when viewed as conical helices, the helical slope, or *lead angle*, of the three-dimensional centerline curve of the shell, is a useful parameter for understanding observed intra– and interspecific morphometric variation (Filin 2026*b*). (See also recently De Renzi & Mayoral 2024, 2026 on helical burrow trace fossils; e.g., ‘devil’s corkscrews’ or *daemonelix*, in keeping with the theme of the title.) Here, I extend this work also to ontogenetic variation.

A large body of recent literature in astronomy is concerned with the shape of spiral arms in galaxies. Especially, how their pitch angle (*ψ* in Fig. 1A) varies among and within galaxies, and within arms (Russell & Roberts 1992; Savchenko & Reshetnikov 2013; Honig & Reid 2015; Pringle & Dobbs 2019). Variation in the ‘tightness’ of spiral arms is discussed in relation to classification and measurable properties of galaxies, such as mass, rotational velocity, and star formation rate (Ringermacher & Mead 2009; Masters *et al*. 2019; Yu & Ho 2019; Sellwood & Masters 2022; Smith *et al*. 2022; Reshetnikov *et al*. 2023). To those familiar with the *Gryphaea* debate of the 1960s and early 70s (Philip 1962, 1967; Burnaby 1965; Hallam 1968; Gould 1972, 1980, 2002; Hallam & Gould 1975; Jones & Gould 1999), as well as earlier and later debates on shell form (Moseley 1838, 1842; Naumann 1840, 1845; Blake 1878; Raup 1966; McGhee 1980; Schindel 1990; Ackerly 1992; Aldridge 1998, 1999; van Osselaer & Grosjean 2000; McGhee 2001), this current literature within astronomy may seem like déjà-vu.

**Figure 1:**
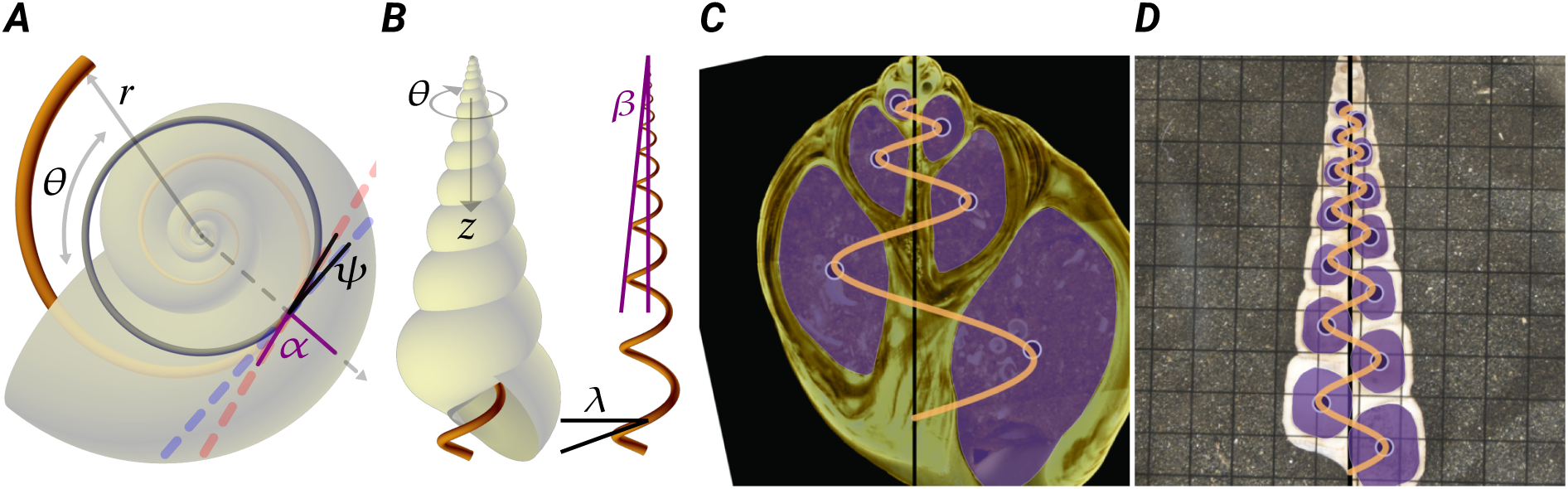
Theoretical morphology and morphometrics of gastropod shells. (***A,B***) Apical and lateral views of shells, defining the various geometric variables and parameters: radius-vector, *r*, winding angle, *θ*, coiling axis (longitudi–nal direction), *z*, spiral and expansion (or pitch) angles for planispirals, *α* and *ψ*, apical semiangle and lead angle for conispirals, *β* and *λ*. The centerline, extracted from the shell in panel B, is a clear example of a conical helix. (***C,D***) Two examples of sectioned shells from the dataset of Collins *et al*. (2021*a*) (C: *Tylospira glomerata*; D: *Maoricolpus roseus*). As part of the data reanalysis, I generated `svg` images, with highlighted apertures (shaded purple), aperture centroids (highlighted circles), coiling axes (central vertical line), and fitted conihelical paths (light orange). The illustrated shells demonstrate the range of shapes and sample sizes (number of apertures/centroids), to which conihelical curves were fitted. More examples in Fig. S1–S3,S6 of SI. Compared to original `jpg` files, `svg` images are zoomed in on specimens and rotated, to obtain vertical coiling axes.

In this paper, I adopt the use of the pitch angle from astronomy, rebranded *expansion angle*. Together, lead and expansion angles fully define the shape of conihelical shell centerlines, with useful applications to mathematical and statistical modeling, to optimality considerations, to morphometric data analyses, and when isometric self-similarity breaks, to quantifying and distinguishing among different forms of allometry. These two centerline parameters also offer baseline expectations for other aspects of shell morphology, such as the inclination of the aperture, i.e., its orientation angles relative to the shell or soft body, and to understanding growth gradients around the aperture, which are one suggested driver of coiling and morphogenesis of gastropod shells (Hammer & Bucher 2005; Urdy *et al*. 2010; Shimizu *et al*. 2013; Johnson *et al*. 2019; Ohta *et al*. 2025). Finally, I re-examine and draw consequences to adaptational explanations of morphological variation in gastropod shells, in light of the constructional-functional-historical triality of phenotypes, as manifested by conihelical geometry.

## Methods

### General and regression formulas for planispiral coiling

The spiral angle, *α*, between the tangent to the spiral and the radius-vector from the spiral’s pole, is a constant for logarithmic spirals. Hence, the alternative name, ‘equiangular’ (Thompson [1942] 1992). Spiral angle is the traditional geometric parameter for describing expansion of planispiral molluscan shells (Moseley 1842; Thompson [1942] 1992). The radius of a logspiral expands exponentially with the *winding angle*, *θ*, such that *r* = *r*_0_*e^γθ^*. The expansion rate, *γ*, is a constant, related to the spiral angle, *γ* = cot *α*. A second way to relate spiral angle to shell expansion is through arclength, ℓ, along the planispiral. Increments of arclength are related to increments of radius by dℓ/d*r* = sec *α* = 1/cos *α* (Philip 1962; Burnaby 1965).

In the study of spiral arms in galaxies, astronomers usually use the pitch angle, instead of spiral angle (Russell & Roberts 1992; Savchenko & Reshetnikov 2013; Sellwood & Masters 2022). To avoid a terminology trap, regarding different meanings of ‘pitch’ (for spirals, for helices, and for general three-dimensional rotations), I will refer to this complement of the spiral angle as *expansion angle*, *ψ* = *π*/2 − *α*. It measures the deflection of the planispiral with respect to a circular path at the same radius (Fig. 1A; a spiral with *ψ* = 0, *α* = *π*/2, follows a circular path). The expansion rate of a logspiral is then given by *γ* = tan *ψ*. A third equivalent definition of spiral angle follows from an often used formula in studies of galactic arms (e.g., Ringermacher & Mead 2009),

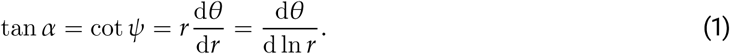

For general planispirals (i.e., non-logspiral), *α* and *ψ* are functions of *r*, changing along the spiral, and consequently also with winding angle, *θ*. In fact, Burnaby (1965, 1966) used a slightly modified version of Eq.[1], in terms of arclength (*r*d*θ* = dℓ cos *ψ*), but limited his analysis to a power-law form of ℓ(*r*). The general solution for any planispiral,

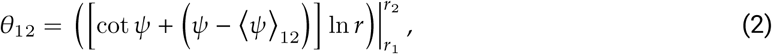

clearly depends on deviation of expansion angle from its (weighted) mean, *ψ* _12_, along the spiral segment from *r*_1_ to *r*_2_ (SI).

While expansion angles for spiral arms of galaxies typically vary between 8^◦^ to 40^◦^ (Pringle & Dobbs 2019; Lingard *et al*. 2021), representing fast-expanding untight spirals, common also to brachiopods and bivalves (e.g., *Gryphaea*), expansion rates for gastropods are usually less than 0.2 (Thompson [1942] 1992; Cameron 1981; *ψ* ≤ 11.3^◦^; Fig. 1A); a consequence of exhibiting several complete revolutions. Taking ln *r* as size measure, the case of a linear-log expansion angle, *ψ* = *a* + *b* ln(*r*/*r*_0_), can be solved exactly (SI),

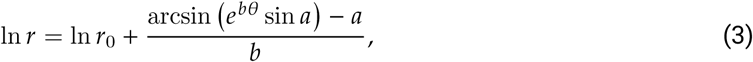

where *r*_0_ = *r*(*θ* = 0) is initial radius, *a* is initial expansion angle, and *b* is its rate of change. This expression becomes the logarithmic spiral, ln *r*_0_ + *θ* tan *a*, in the limit *b* → 0; and, if *ψ* remains small throughout ontogeny, reduces to a more user-friendly form, ln *r*_0_ + (*a*/*b*) *e^bθ^* − 1 (SI). Either form can be used as an alternative to quadratic regression of ln *r* on *θ* (McGhee 1980; Schindel 1990; Ackerly 1992), when testing for deviations from perfectly logspiral coiling.

### Optimal planispiral growth

An interval of planispiral growth is defined by an increase in radius from *r*_1_ to *r*_2_ over a winding angle span *θ*_12_. The shortest path from (*r*_1_, 0) to (*r*_2_, *θ*_12_) is obviously the straight line between them. From an optimal control perspective, the curve defining a straight line is the *singular arc*, in which the value of the control variable, here expansion angle *ψ*, can vary freely, to obtain the optimal path. In euclidean space, singular control translates to straight lines.

However, if *ψ* is constrained to vary only between 0 and *M* (*M* = const > 0), the optimal path from (*r*_1_, 0) to (*r*_2_, *θ*_12_) can no longer follow a straight line. Instead, the optimal path will be a combination of circular paths, where expansion angle is fixed at its lower boundary (*ψ* = 0), straight line segments (i.e., episodes of singular control, in which *ψ* varies freely inside the interval [0, *M*]), and logspiral segments of maximum expansion, *ψ* = *M* (i.e., expansion angle fixed at upper boundary). (As also obtained in optimal control studies in robotics, Dubins 1957; Reeds & Shepp 1990; Bhattacharya *et al*. 2007; the last study referring to such line and logspiral segments [including circular paths] as “the language of shortest paths”). A circular path is clearly minimized when its radius is minimized, i.e. at the initial radius *r*_1_. On straight lines, expansion angle increases with winding angle (given *ψ* ≥ 0). Consequently, we can conclude that optimal planispiral growth, by a ratio *q* = *r*_2_/*r*_1_ ≥ 1 over a winding angle span *θ*_12_, is composed of an initial circular path (*ψ* = 0), followed by a straight-line segment (*ψ* growing from 0 to *M*), and finally terminating with a logspiral phase (*ψ* = *M*). A more detailed optimal control analysis is provided in SI.

Clearly, this result can receive an adaptational interpretation, of obtaining a given radial growth ratio, *q*, with minimum investment in arclength growth. It also has a more immediate application, for data analysis, as it provides the minimum arclength between two points. Both these considerations are discussed further in Results and Discussion.

### Conispiral (conihelical) coiling

Non-planispiral shells are obtained by adding to a planispiral, ℓ(*θ*), some translation along the coiling axis, *z*-direction, such that each increment of planispiral (arc)length, dℓ, is associated with a perpendicular increment, d*z*. If the increments are always proportional to each other, d*z* ∝ dℓ, the resulting spiral space curve is defined by a fixed slope angle, *λ*, termed *lead angle*, d*z* = dℓ tan *λ*. Such space curves are called *generalized helices*, or *curves of constant slope* (Nutbourne & Martin 1988; Scofield 1995), of which the familiar circular helix (springs and corkscrews) is a special case (ℓ (*θ*) a circle, *ψ* = 0).

For a logarithmic planispiral, the corresponding space curve is a *conical helix*, winding around a right circular cone, with apical semiangle *β* (Fig. 1B). This is the logarithmic *conispiral* of Moseley (1842), Thompson ([1942] 1992) and Raup (1966). Raup’s translation parameter is *T* = cot *β*. Expansion rate is now *γ* = cot *α* sin *β*, where spiral angle, *α*, is measured on the conical envelope, rather than on the corresponding planispiral (Moseley 1842; Thompson [1942] 1992; Raup & Graus 1972; Løvtrup & von Sydow 1974; Ekaratne & Crisp 1983). Consequently, the relation *α* + *ψ* = *π*/2 holds only for planispiral coiling, and expansion angle is now given by tan *ψ* = *γ* = cot *α* sin *β*. Raup’s well-known *whorl expansion rate* parameter, *W*, is an equivalent way to define expansion, *W* = *e*^2^*^πγ^* or *γ* = ln *W*/2*π*.

Lead angle, apical semiangle, and expansion angle are interdetermined through

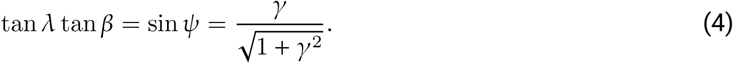

Given *γ* ≤ 0.2 for most gastropods, this equation can be approximated by tan *λ* tan *β* ≈ *γ* (see SI). A version of Eq.[4] was already presented by Naumann (1840), in terms of *W*, rather than *γ* or *ψ*. As briefly reviewed by Raup (1966), subsequent work however gravitated toward planispiral coiling in ammonoids and kin, and treatment of conispirals was continued by Moseley (1842), and later by Blake (1878).

### Allometric coiling

The shape of a conihelical *centerline* of a shell is wholly defined by only two parameters. In this study, I opt for lead and expansion angles, *λ* and *ψ*, which then determine the other parameters (*α*, *β*, *γ*, *W*, *T*) through the above formulas. A consequence of this high integration and tight covariation of coiling parameters and morphometric traits is that it can break in many different ways. Allometry thus may be obtained through different geometries.

First, allometric generalized helices are defined by a constant lead angle, but varying expansion angle. This is the most commonly observed form of allometry in studies that apply Raup’s *W*-*T* morphospace (Davoli & Russo 1974; Newkirk & Doyle 1975; Ekaratne & Crisp 1983) or other conispiral parametrization (van Osselaer & Grosjean 2000). Coiling rate, or tightness (Joysey 1959; Philip 1962, 1967; Burnaby 1965), varies around the fixed coiling axis. Because the helical slope, tan *λ*, is still constant, there is still simple proportionality between centerline height, *z*, and planispiral arclength, ℓ, Centerline radius, *r*, on the other hand, is not proportional to ℓ, as the base planispiral, ℓ(*θ*), is no longer logarithmic. Consequently, some common allometric relationship for ℓ(*r*) and *z*(*r*) applies (e.g, Burnaby’s power-law). Specific cases of such generalized helical shells (e.g., for arithmetic planispirals; Fig. 5A) were considered by Rice (1998) and Urdy *et al*. (2010).

An obvious second way to break conispiral isometry is to fix expansion angle but vary lead angle. In such cases, the corresponding base planispiral, ℓ(*θ*), is logarithmic, and therefore ℓ ∝ *r*. Centerline height, *z*, however, is no longer proportional to ℓ (or to *r*). A special case of this type of allometry is the logarithmic helicospiral model, where both *r* and *z* expand exponentially, but with different expansion rates, *γ_z_* ≠ *γ_r_* (Kohn & Riggs 1975; Bayer 1978; Schindel 1990; Savazzi 1990; Fowler *et al*. 1992; Stone 1995).

More complex allometries, where the coiling axis itself may change, can be studied with differential geometry or stereography (Okamoto 1988, 1996; Ackerly 1989*b*; Moulton *et al*. 2012; Moulton & Goriely 2014; Filin 2026*b*). However, if change in lead and/or expansion angle is small, shell centerlines are still fairly regular, albeit no longer perfectly self-similar. Finally, allometry can also manifest in other aspects of the growing ‘tube’ (Okamoto 1988), or ‘helicocone’ (Cox 1960), that is the coiled shell. Particularly, in relative size and shape of the aperture, or whorl cross-section.

### Datasets

I previously (Filin 2026*b*) compiled reports of population and sample means of coiling parameters from Newkirk & Doyle (1975), for *Littorina saxatilis*, and Davoli & Russo (1974), for fossil and recent *Subula* (syn. *Terebra*), as well as data for over 400 extant land, freshwater and marine gastropod species from Noshita *et al*. (2012) and Araki & Noshita (2023), based on single museum specimens. Here, I reuse this previously digitized dataset (Filin 2026*a*).

In this study, I additionally include Collins *et al*. (2021*a*), containing ontogenetic sequences of apertures of fossil and recent marine gastropods of the New Zealand region, obtained from sectioned specimens or microtomography. This dataset contains both images of sectioned specimens and morphometric measurements of coiling axes, aperture areas, and aperture centroids (Fig. 1C,D, Fig. S1–S3,S6), thus providing opportunity to directly estimate spiral centerlines, their potential allometry, and the separate allometry of apertures.

### Data reanalysis

Values of expansion rate, *γ*, apical semiangle, *β*, lead angle, *λ*, spiral angle, *α*, and expansion angle, *ψ*, can be obtained from reported values of Raup’s *W* and *T*, given the above formulas. Downward and outward inclination angles of shell apertures, Δ and Γ respectively, from Noshita *et al*. (2012) and Araki & Noshita (2023), were also reanalyzed (see below, Results and Discussion and Fig. 3). I reversed the sign of Γ, for consistency with expansion angle, *ψ* (i.e., outward inclination of either centerline or aperture is positive; inward negative).

For the Collins *et al*. dataset, base planispirals were fitted with ln *r* ∼ *θ* linear regression, both OLS and major-axis model II (hereafter, MA). Quadratic and nonlinear regressions were also performed to test for ontogenetic variation in expansion angle (e.g., Eq.[3]). I also applied model selection, to compare various per-specimen *r*(*θ*) models, by adapting and enhancing code from Collins *et al*. (2021*a*).

In that dataset, longitudinal coordinate, *z* (Fig. 1B), is measured with respect to a specimen-dependent arbitrary datum, taken to be the first (i.e., youngest) aperture in the ontogenetic sequence (Fig. 1C,D). Denoting this measure of spire height by *y* (as in Collins *et al*. 2021*a*, *b*), ontogenetic sequences of aperture centroids always start at *y* = 0. Number of sampled apertures, however, varies markedly among specimens, even within the same species (Fig. 1; Fig. S1, S3, S6). In other words, the origin of longitudinal measurement is specimen-specific and not standard among all specimens and species. Collins *et al*. (2021*b*) correctly pointed out that, in such cases, there is a need for an intercept parameter that “accounts for the variation in starting point”. This, again, alludes back to the *Gryphaea* debate, and to Gould’s (1972) correction of Burnaby’s (1965) mistake of comparing intercepts, rather than the full ontogenetic trajectories. However, Collins *et al*. later fit linear models to log-transformed response variables (their ‘Exponential’, i.e. logspiral, and ‘Power law’ models), where intercepts no longer provide the needed correction.

Here, I circumvent this “insidious… pitfall that could lead to confusion with allometry” (van Osselaer & Grosjean 2000), by applying the expected linear relationship *y* ∼ ℓ for conical helices, where both *y* and ℓ are measured relative to the specimen-specific datum. I apply to *y*(ℓ) the same model selection, and linear and quadratic regressions, such that ontogenetic variation in lead angle can also be detected. In addition, I estimated the apical semiangle, *β*, for Fig. 2, by *y* ∼ *r* linear regression.

**Figure 2:**
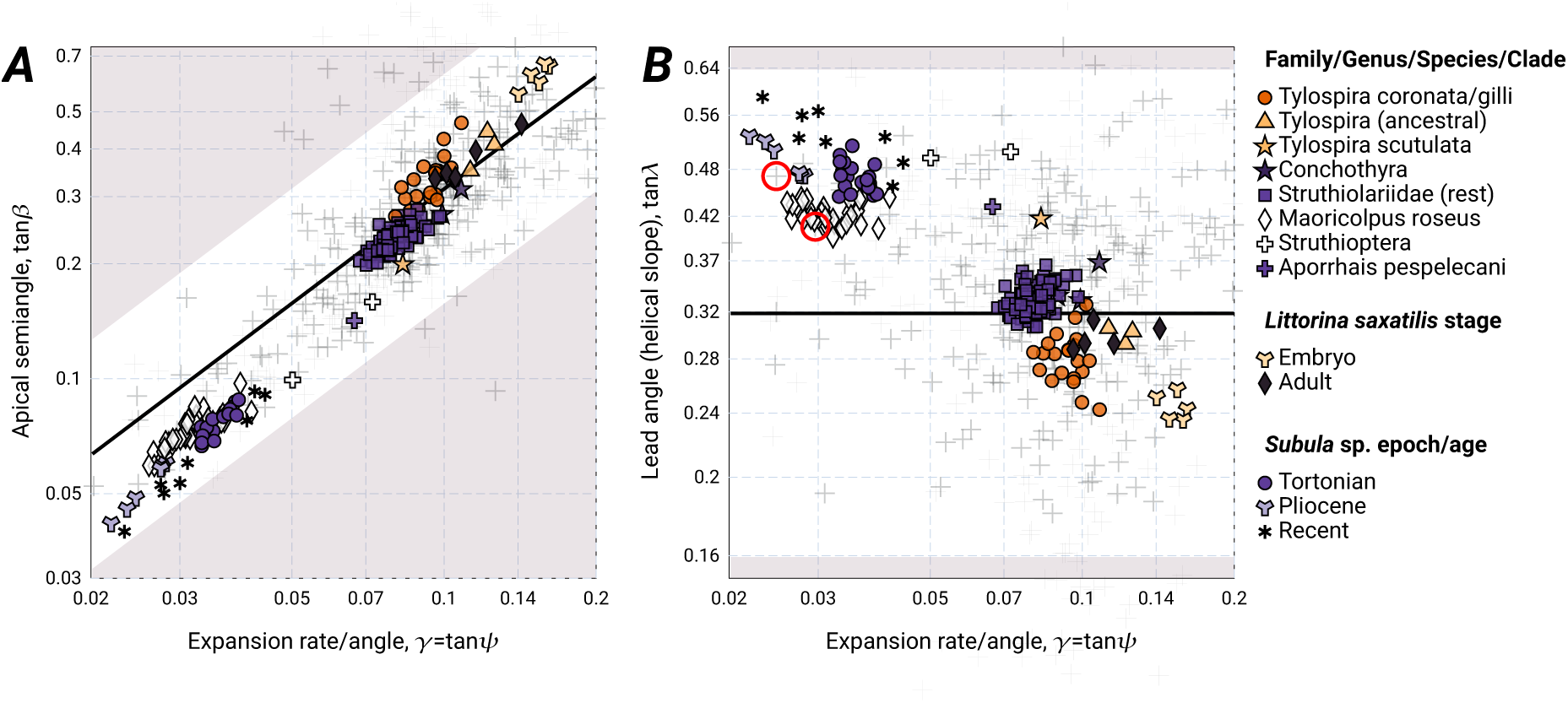
Covariation of coiling parameters in the reanalyzed datasets. (***A***) Relationship between apical semiangle, tan *β*, and expansion rate, *γ* (= tan *ψ*). Marine species from Noshita *et al*.’s data are presented in gray, as background scatter and reference of the total variation exhibited in this group. tan *β*-values for Collins *et al*.’s data are from *y* ∼ *r* regressions, as described in Methods. Taxa presented are the Struthiolariidae (including separately the *Tylospira* clade, which is further divided into later forms and to ancestral forms, *Perissodonta* and *T. glomerata*), *Maoricolpus roseus* (Turritellidae), and the three Aporrhaidae specimens of *Struthioptera camachoi*, *S. haastiana* and *Aporrhais pespelecani*. For all other datasets (population means of *L. saxatilis* from Newkirk & Doyle, sample means of *Subula* species from Davoli & Russo, and data of Noshita *et al*. and Araki & Noshita), tan *β*-values were obtained from reported values of Raup’s *T* (tan *β* = 1/*T*; SI). The unshaded area corresponds to 1/2*π* ≤ *λ* ≤ 2/*π* (Filin 2026*b*), where most of the data lies. The solid black line represents *λ* = 1/*π*. Medium-spired shells (e.g., *Littorina* and Struthiolariidae, including *Tylospira*) seem to fall close to this line. Both within groups and overall, there is a clear direct relationship between apical semiangle and expansion rate. This tight correlation is explained by (***B***) variation in growth (expansion rate) coupled with more-or-less fixed lead angles within each clade or sample. Values of tan *λ* for Collins *et al*.’s data were estimated from *y* ∼ ℓ regressions. For all other datasets, tan *λ*-values were obtained from reported values of Raup’s *W* and *T*, using formulas presented in Methods (e.g., Eq.[4]). Circled red are specimens of *Turritella rubescens* and *T. fortilirata* from the Noshita *et al*. dataset. These coincide well with the scatter for *Maoricolpus*, thus confirming the validity of the two methods of obtaining tan *λ*-values. A mixed-effect linear growth model estimated tan *λ* at 0.421 [0.417, 0.425] for *Maoricolpus*, 0.283 [0.277, 0.288] for the *Tylospira* clade, and 0.330 [0.327, 0.333] for rest of the Struthiolariidae.

All data analyses were done in `R` (version 4.3.3; R Core Team 2024), applying various packages (Warton *et al*. 2012; Legendre 2024; Elzhov *et al*. 2023; Pinheiro *et al*. 2025; Bates *et al*. 2015; Lüdecke *et al*. 2021; Kuznetsova *et al*. 2017; Stoffel *et al*. 2017; Wickham & Henry 2023; Wickham *et al*. 2023; Mazerolle 2023). I also generated graphically annotated `svg` images of specimens, for convenient visual inspection of data and fitted conihelical curves (Fig. 1C,D; Fig. S1–S3,S6). More details and results on data analyses in SI. All data, code, and images are available in online repositories.

### Mixed-effect growth models

Testing for inter-clade differences, while accounting for individual variability in growth and allometry, was accomplished with mixed-effect linear growth models (Grimm *et al*. 2017). The three main clades of the Collins *et al*. dataset are: *Maoricolpus roseus* (Turritellidae), *Tylospira* +*Perissodonta* (mostly *Tylospira coronata*), and the rest of the Struthiolariidae (mainly *Pelicaria vermis* and *Struthiolaria papulosa* specimens). (34, 25, and 88 specimens respectively, out of total 164.) I estimated and tested for differences in allometry and coiling parameters. In particular, in the growth and allometry of aperture size, *A*, with respect to other morphometric measures (*r*, *θ*, ℓ, etc.).

### Measurement error, misaligned coiling axes, and regression weighting

Repeatability of coiling parameter estimates, within and among specimens, was investigated (*i*) by using the repeatedly resampled/resectioned specimens in the Collins *et al*. dataset, originally used to estimate measurement (digitization) error; (*ii*) by correcting for potentially misaligned coiling axes in the original data, based on left-right bisector of centroids (e.g., Davoli & Russo 1974); and (*iii*) by exploring different weighting for centroids in within-specimen regressions (either no weighting, or weights by aperture size).

Nevertheless, realigned coiling axes, in the vast majority of specimens, were practically indistinguishable from the manually measured axes (Fig. S3 in SI). Ultimately, uncertainty in parameter estimates due to sampling (digitization) and axis misalignment was practically negligible, and due to regression weighting was still relatively small. Specimen-level within-group repeatabilities of expansion rate were 0.935 [0.877, 0.962] (95%CI), 0.827 [0.764, 0.869], and 0.798 [0.67, 0.873], for the *Tylospira* clade, rest of Sturthiolariidae, and *Maoricolpus*, respectively. Estimates of helical slopes, tan *λ*-estimates, exhibited even higher repeatabilities (respectively, 0.972 [0.942, 0.985], 0.892 [0.847, 0.921], 0.974 [0.955, 0.984]). Total specimen repeatabilities, including group differences, were 0.986 [0.982, 0.989] for expansion, and 0.993 [0.991, 0.995] for lead (more details in SI).

Beyond minor shifts in numerical values, all results and conclusions of this study remain unchanged, regardless of weighting or coiling-axes realignment (e.g., Fig. S4–S5). I therefore present in the main text results based on originally (manually, visually) measured coiling axes (Collins *et al*. 2021*a*) and with no weighting.

## Results and Discussion

### Optimal control in planispiral growth

When studying optimal growth paths over half revolutions, i.e., *θ*_12_ = *π*, as in successive whorl cross-sections of sectioned shells (Fig. 1C,D), the shortest arclength, for a given diameter *r*_1_ + *r*_2_ = *d* = const, is 1.4573*d*, if maximal expansion angle is *M* = arctan(0.2). This shortest path occurs at *q* = 1/W_0_ *e*^(^*^M^*^−^*^π^*^)^ ^tan^ *^M^*^+sec^ *^M^* cos *M*, which is 1.373 in this specific case, i.e., radial growth by 37% through a half revolution (equivalent to *γ* ≈ 0.1; W_0_ is the zeroth branch of the Lambert W function; not to be confused with Raup’s *W* coiling parameter).

The maximum path is defined by a semicircle, of arclength *πd*/2, at *q* = 1 (i.e., no radial growth; *π*/2 = 1.5708). In other words, the maximum relative variation in arclength is 7.5%, if expansion rates are constrained between 0 and 0.2. When comparing only fully logspiral paths, the expression for arclength is tanh(*πγ*/2) csc *ψ d*, and relative variation in arclength reduces to 1.3%.

The immediate practical consequence of these results is that, for gastropods, the approximation *πd*/2 for arclength between successive whorl cross-sections is a very reasonable one (maximum 7.5% error; or 1.3% if assuming that growth is by piecewise logspiral increments). This result has application value in data analyses, for estimating arclength from photographs.

The adaptational consequence is that, for gastropods, deviation from the optimal growth path results in wastage (i.e., extra arclength) of only few percent at most, and that logspiral paths are fairly close to optimal. In other words, such a selective force is likely to be overwhelmed by other considerations, adaptational or not, that may affect shell morphology. Such economical shortest growth path over a given winding angle span, is likely to play a more significant role when expansion angles are higher, as in bivalves and brachiopods. A similar argument was made by Hutchinson (1990), in relation to control of global shell shape by apertural growth gradients; i.e., more likely in bivalves than in gastropods.

### Fit of conihelical and allometric coiling

Model selection consistently preferred exponential *r*(*θ*) and linear *y*(ℓ), as expected for conical helices (Table S1). Per-specimen visual inspection of fitted conihelical curves confirmed the very good fit in the vast majority of cases (Fig. 1C,D; Fig. S1–S3,S6).

Limiting the discussion hereafter to the three main clades of the Collins *et al*. dataset (148 specimens out of 164), linear regression of ln *r* ∼ *θ* produced very good fits (*R*^2^ ≥0.9991 for MA; *R*^2^ ≥0.9 for OLS, except for two specimens [0.84 and 0.894], due to having only 4 centroid points and slight coiling-axis misalignment; Fig. S3*N,P*). Expansion rates from OLS and from MA are practically identical (*r*^2^>0.9999). Similar results were obtained for *y* ∼ ℓ linear regressions (*R*^2^ ≥0.9986; *R*^2^ ≥0.9737; *r*^2^ >0.9999).

Of 148 specimens, 89 did not exhibit any significant trend in expansion or lead angle. Of the remaining 59, 41 showed a trend in expansion angle, based on quadratic OLS or Eq.[3] (*p* ≤ 0.05), and 30 in lead angle (quadratic OLS), with 12 specimens having a significant trend in both lead and expansion. Applying Bonferroni-Holm correction within each of four groups (*Tylospira* clade, *Pelicaria*, rest of Struthiolariidae, and *Maoricolpus*), these counts decreased respectively to 9, 4, and 1. Except for three cases (out of 41) identified exclusively by Eq.[3], and two cases identified exclusively by quadratic OLS, both regression methods agreed on the same 36 cases of change in expansion angle, and on all nine specimens after *p*-value adjustment.

More importantly, when significant, the median per-specimen relative change in lead angle was 26%, i.e., roughly ±13% around the linear regression estimate. For expansion angle, median relative change was roughly ±33%. Change in expansion was almost exclusively negative (*b* < 0 in Eq.[3]), with faster expansion in the smaller younger whorls, as also observed in previous studies (Davoli & Russo 1974; Newkirk & Doyle 1975; Vermeij 2020).

Model selection identified a power-law *A*(*r*) as the preferred model for aperture size. Isometric self-similarity dictates *A* ∝ *r*^2^. Mixed-effect comparison among clades reveals that aperture area grows with slight positive allometry for *Maoricolpus* (allometric exponent is 2.11, 95%CI [2.07, 2.15]), but with negative allometry for Sturthiolariidae — 1.73 [1.66, 1.79] for the *Tylospira* clade, and 1.86 [1.83, 1.89] for the rest of Struthiolariidae. The larger deviation from an exponent of 2 for *Tylospira* reflects also the wider and faster-expanding columella in this clade (clearly visible in Fig. 1C). Vermeij (2020) recently discussed skeletal dynamism, in the context of inner whorl resorption and shell remodelling. Breaking of self-similarity, through growth allometry of the helicocone cross section, relative to its length extension (centerline arclength), may be a more common way of overcoming constraints of (log)spiral growth. A path of ‘least resistance’ for achieving a rapid change in overall morphology. Thick columellae result as ‘spandrels’, caused by concentrated shell accretion in the limited umbilical gap that inevitably forms.

Overall, centerline allometry is detected in only a minority of specimens, manifests sporadically among and within groups, and visually barely perceptible or weak in most cases (Fig. S6 in SI). The only consistently strong allometry is that of aperture size in the *Tylospira* clade. Isometric conical helices account for most of the variation within and among specimens and groups (Fig. 2).

Finally, despite their conclusions, I note that Collins *et al*.’s (2021*b*) results also hint at conihelical geometry. In their Table 1, ‘Exponential’ (i.e., logspiral) models fitted aperture size and columellar width relatively well, but performed poorly for *y*-measurements. That was an indication of a potential problem, as morphometric traits must ultimately combine to describe three-dimensional wholes in a consistent way. To quote Thompson, “how… the whole organism hangs together, and how throughout its fabric one part is related and fitted to another… is sometimes apt to be forgotten in the course of…more and more minute analysis by which… we seek to unravel the intricacies of a complex organism… The skeleton begins as a continuum, and a continuum it remains all life long.” (Thompson [1942] 1992).

### Adaptational, constructional, and historical explanations of covariation

Observed covariation in coiling parameters, and uneven distributions of taxa within morphospaces often receive adaptational interpretations, due to mechanical, economical, hydrodynamic, and similar considerations (Raup 1966; Linsley 1977; Noshita *et al*. 2012; Okajima & Chiba 2013; Okabe & Yoshimura 2017; Araki & Noshita 2023; Páll-Gergely *et al*. 2024). However, as discussed in Methods, conihelical morphology exhibits high levels of integration. Covariation between two traits may simply be a passive consequence of growth-related variation in one of them, or in a third trait. In particular, Eq.[4] and Fig. 2 demonstrate that a fixed lead angle, coupled with plasticity in growth (expansion), prescribes a tight relationship between apical semiangle and expansion rate, as commonly observed. Raup’s (1966) famous whorl-overlap condition, which also dictates a relationship between expansion rate and apical semiangle, is practically indistinguishable from Eq.[4], given the low expansion angles of most gastropods (Filin 2026*b*).

Approaching shell morphology through the apical (semi)angle may be a legacy of the verbal conchological classification of shells (i.e., turreted, turbinate, discoid, etc.). However, it may miss the constructional and developmentally plastic aspects that dominate at lower taxonomic and the organismal levels, as prescribed by Eq.[4] and shown in Fig. 2. Adopting *ψ* and *λ* as the natural axes of coiling behavior is also strongly supported by experiments and observations on within-species developmental plasticity, such as scalariforms and “corkscrew”s (Hutchinson 1989; Checa *et al*. 1998; Clewing *et al*. 2015), and effects of variation in soft-body growth on shell morphology (Vermeij 1980; Kemp & Bertness 1984; van der Sman *et al*. 2009).

Such potential constructional effects are also evident when considering the inclination angles of apertures. The angles *ψ* and *λ* measure the orientation of the growing tip of the centerline, with respect to a cylindrical coordinate system (Fig. 1). Similarly, outward and downward aperture inclinations, Γ and Δ (Noshita *et al*. 2012), measure the orientation of the aperture — the opening of the shell through which “a snail must conduct all of its business with the outside world” (McNair *et al*. 1981). When these two orientations perfectly match, i.e., Γ = *ψ* and Δ = *λ*, the shell’s aperture is perpendicular to the conihelical centerline (’orthoclinal’ *sensu* Cox 1960; Illert 1989). Γ = 0 defines a ‘radial’ aperture, *sensu* Linsley (1977), and Γ > *ψ* and/or Δ > *λ* characterize a ‘tangent’ aperture.

The deviations, or *relative inclinations*, Δ^∗^ = Δ − *λ* and Γ^∗^ = Γ − *ψ*, may therefore be better indicators of constraint and/or adaptation (Okajima & Chiba 2013). As Fig. 3B demonstrates, relative inclinations are roughly centered around (0, 0), for 428 species of marine, freshwater, and land gastropods. The three habitat groups, however, are distributed differently in (Γ^∗^, Δ^∗^)-space, with excess of Δ^∗^ < 0 for freshwater, excess of Γ^∗^ > 0 for marine, and excess of Γ^∗^, Δ^∗^ > 0 for land snails; the last being practically absent from the opposite quadrant (Fig. 3B; SI).

**Figure 3:**
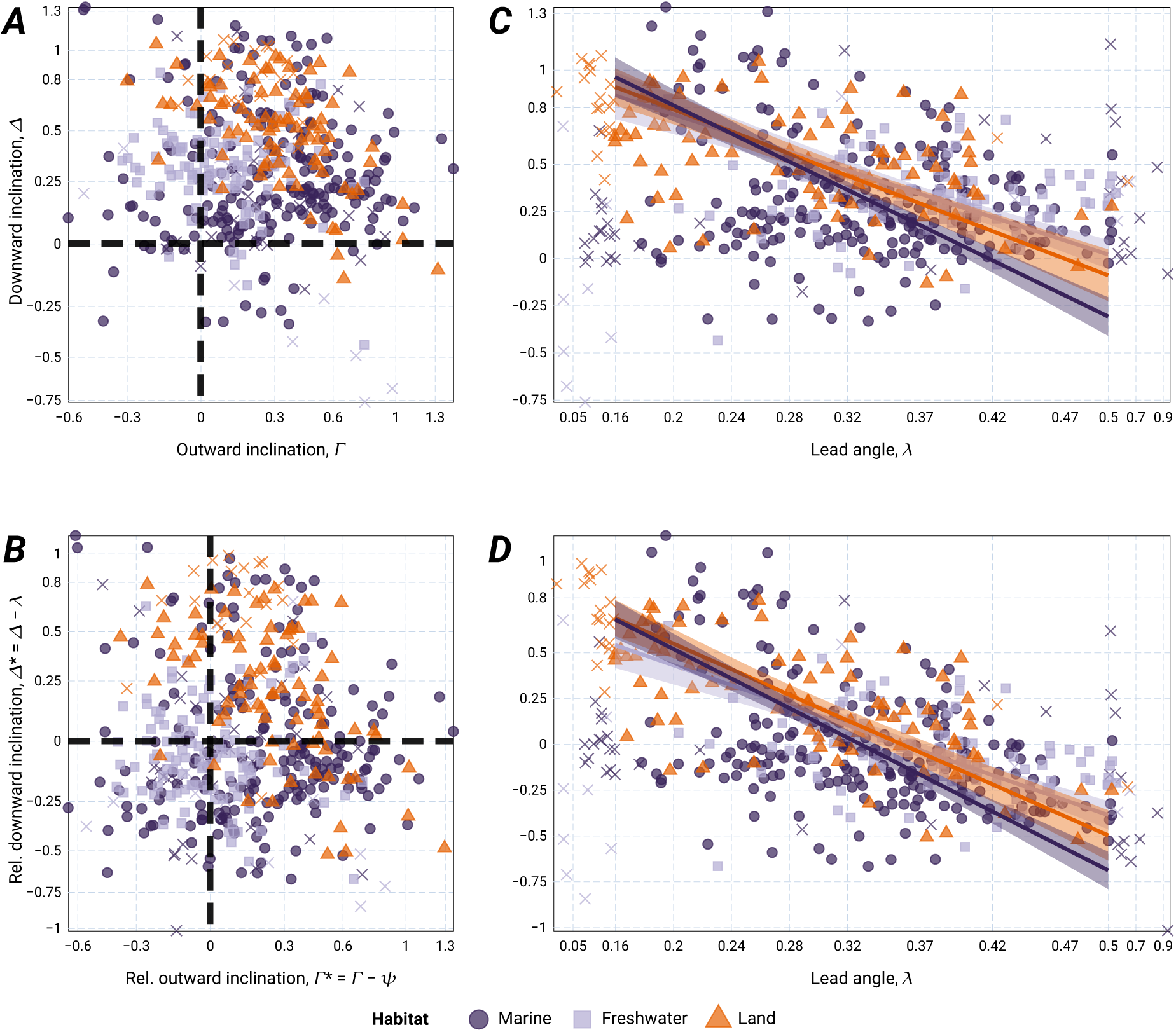
Outward and downward aperture inclinations. Specimens outside the ‘main sequence’ (see main text) are marked with ×. (***A***) There is a clear bias toward both positive downward (Δ) and positive outward (Γ) inclinations. That is expected, given that centerline direction, *λ* and *ψ*, sets a baseline orientation for the shell and soft body. Deviations from this baseline are measured by (***B***) the relative inclinations, Δ^∗^ and Γ^∗^. As discussed in main text and SI, the three habitat groups differ in distribution of relative inclinations. Both (***C***) (absolute) downward inclination, Δ, and (***D***) relative downward inclination, Δ^∗^, tend to decrease as lead angle, *λ* increases. For freshwater species, the null hypothesis of uncorrelated Δ (but not Δ^∗^) and *λ* could not be rejected. Per-habitat model II SMA line fits are presented, with corresponding 95% confidence intervals. The *λ*-axis is piecewise linear (80% of the scale is between 0.16 and 0.5), to highlight the variation in the main sequence.

Borrowing again from astronomy (as did McNair *et al*. 1981), a *main sequence* of gastropod coiled morphology can be defined by 0 ≤ tan *ψ* ≤ 0.2 and 0.16 ≤ tan *λ* ≤ 0.64 (Methods; Fig. 2; Filin 2026*b*). Within this majority of observed gastropod morphologies (369 species), I applied OLS, SMA model II, and mixed-effect linear regressions to explore patterns of covariation between aperture inclination and centerline coiling, while controlling for habitat as a fixed effect, and for taxonomic family as random intercepts. All methods supported the same conclusions, and patterns were robust under several resampling and bootstrapping procedures, including removal of small-sample-size families and/or downsampling to even per-family sample sizes (SI).

Freshwater species do not show any trend in aperture inclination, seemingly randomly distributed around Γ^∗^ = 0, Δ^∗^ = 0, or Δ = 0.32 ≈ 1/*π* (Fig. 3A,B). Land and marine species share the same inverse relation between Δ or Δ^∗^ and lead angle, *λ* (i.e., common slope). Thus, rejecting the null expectation of a slope of +1 for Δ(*λ*), and 0 for Δ^∗^(*λ*), if downward aperture inclination is distributed randomly around that of the centerline. However, once effects of taxonomic family are considered (random intercepts), this trend is significant only in land snails, having been partially incorporated into inter-family differences (Fig. 4A,B). Moreover, controlling for family effects also in lead angle results in a common inverse relationship between Δ^∗^– and *λ*-residuals for all three habitat realms (Fig. 4C,D; SI).

**Figure 4:**
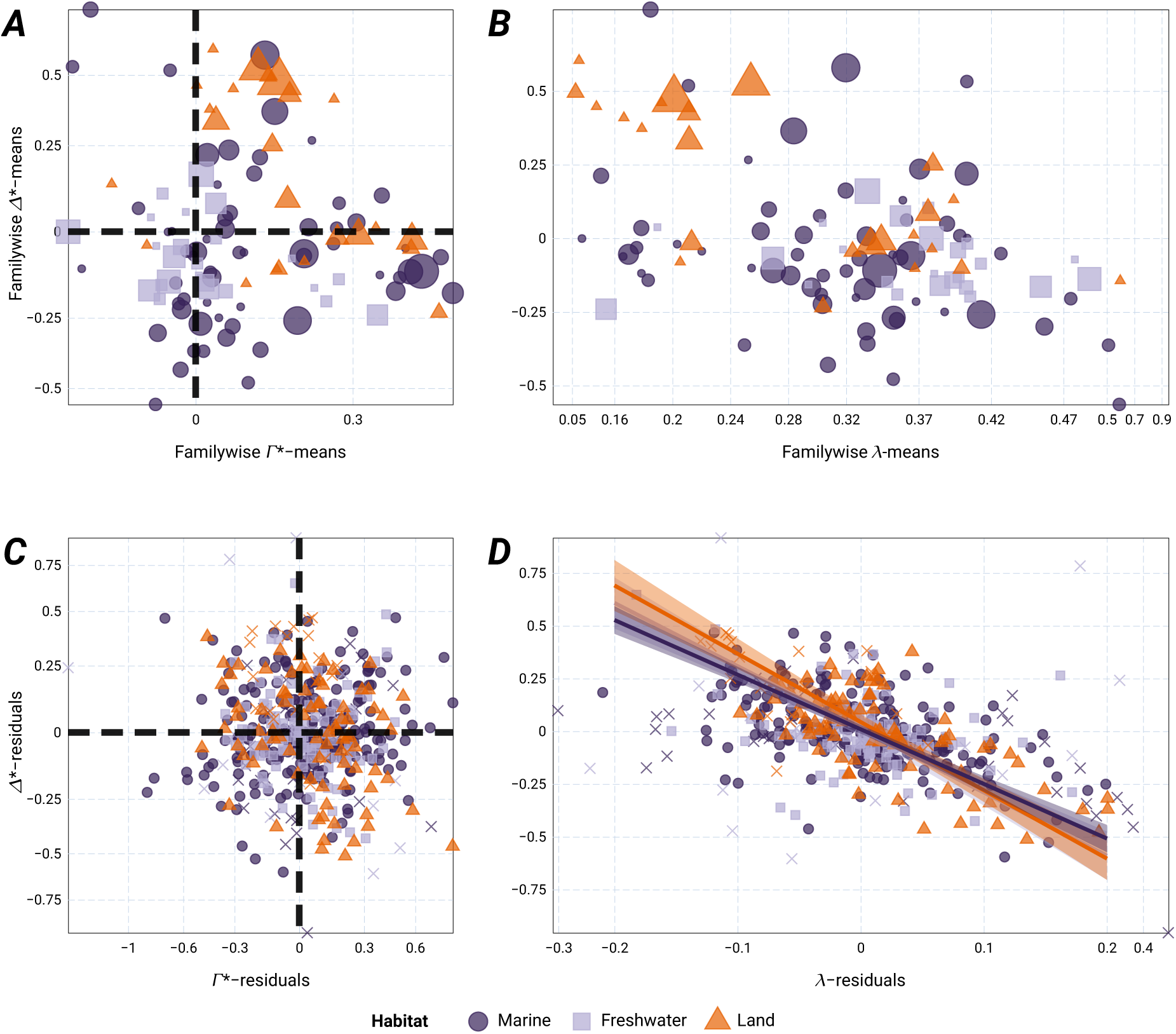
(***A***) Per-family means of relative downward and outward inclinations, Δ^∗^ and Γ^∗^. Symbol (2D) size is a linear function of family size (number of specimens/species in the dataset), ranging from 1 to 32, with mean size 4.4, and median size of 2. (***B***) Relationship between per-family mean downward inclination and per-family mean lead angle; obtained from a random-effect model, with no fixed intercept and taxonomic family as the grouping factor. *λ*-axis piecewise linear (80% of the scale is between 0.16 and 0.5). In both panels the familiar bimodal distribution for spire-height in land snails is visible. Symbol size describes number of dataset specimens belonging to a family; smallest being 1, second smallest being 2, and subsequently linearly increasing to a maximum of 32 specimens (the marine Muricidae). (***C***) distributions of residuals after removing effects of taxonomic family. (***D***) within-family residual lead angle and downward inclination are strongly negatively correlated. The relationship is identical among all three habitat groups (i.e, no significant differences in slope or intercept/elevation). *λ*-axis is, again, piecewise linear (80% of the scale is between –0.2 and +0.2). Specimens outside the ‘main sequence’ (see main text) are marked with ×.

For outward inclination, Γ or Γ^∗^, the only robust pattern is potentially a negative relationship between outward inclination and expansion angle, *ψ*, only in land snails. However, this effect disappears after accounting for familywise random intercepts (Fig. 4A–C; SI).

Outside the main sequence, shells either expand fast (*γ* > 0.2; mostly marine, e.g., *Haliotis*) or are very low-spired/planispiral (*λ* < 0.16). The distribution pattern in Fig. 3C,D suggests that, for the latter group, land, marine and freshwater snails adopt different solutions, in terms of aperture inclination. However, most such freshwater and land ‘planispirals’ in this dataset are of only two families (respectively, Planorbidae and Bradybaenidae).

To summarize, the aperture is an interface between the environment and the shell. Mean values of Γ^∗^ and Δ^∗^ that are either close to zero or positive are understandable, given that, on average, the “outside world” is located outward and downward relative to a shell’s centerline. With faster expansion and steeper helical slopes, the centerline is already comparatively pointing away from the shell, so less correction in aperture orientation is required for contact with the environment (e.g., substrate), resulting in an inverse relationship between Δ^∗^ and *λ* (Fig. 3D; Fig. 4B,D). For high-spired ‘shell draggers’ (McNair *et al*. 1981), the centerline may in fact ‘overshoot’ the substrate, leading to upwards correction (negative Δ^∗^; see also Okajima & Chiba 2013). These are constructional patterns par excellence, observed in most or all habitat realms (Fig. 3; Fig. 4D).

These findings do not preclude additional adaptational effects of centerline orientation and/or relative aperture inclination, such as for defense, mechanical strength, economical shell construction, postural stability, aestivation, or particular ecologies (Raup 1966; Linsley 1977; McNair *et al*. 1981; Noshita *et al*. 2012; Okajima & Chiba 2013; Araki & Noshita 2023; Páll-Gergely *et al*. 2024). However, geometry (centerline orientation with respect to substrate; Vermeij 1971), history (here, effect of taxonomic family; Fig. 4A,B), and chance, all exert their influences in producing uneven distributions within morphospaces, complicating demonstrations of adaptation (Raup & Gould 1974; Pie & Weitz 2005).

### Consequences to morphogenetic explanations

In self-similar isometrically growing conispiral (conihelical) shells, the different spirals that make up the shell’s surface, by geometric necessity, differ in spiral angle, *α*, apical semiangle, *β*, and lead angle, *λ* (Fig. 5B,C). However, they all share a single common value of expansion rate (Illert 1983), *γ* = tan *ψ*.

**Figure 5:**
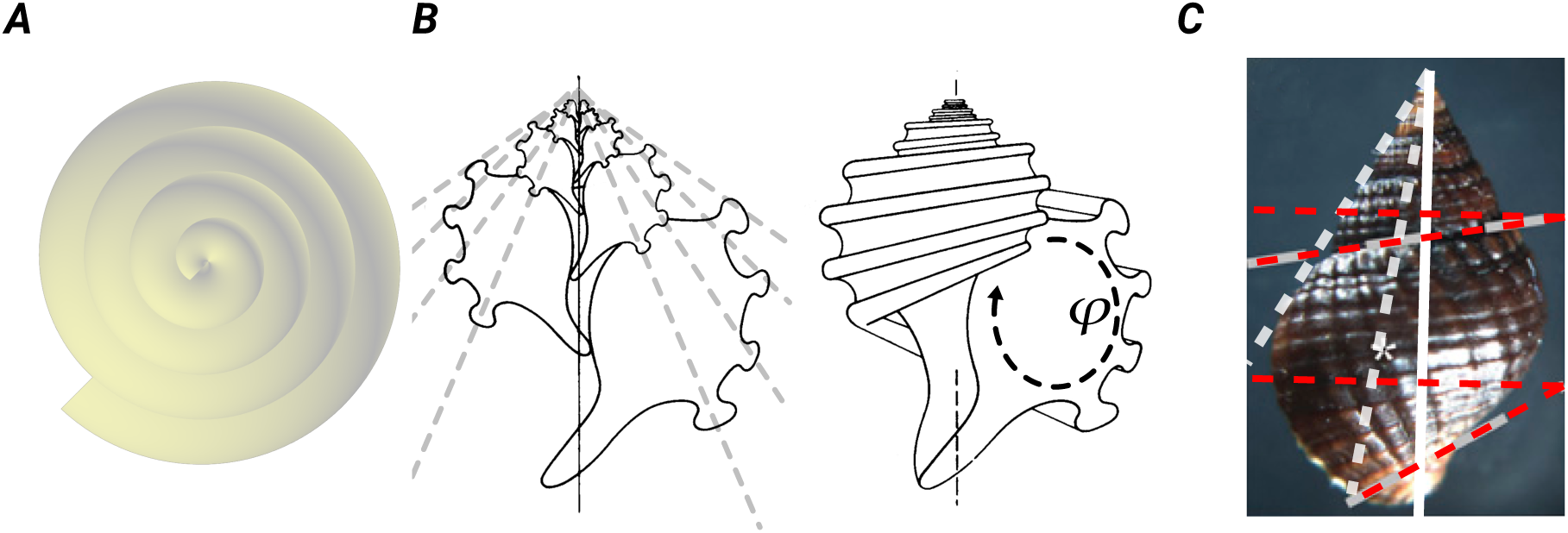
More theoretical morphology in a snailshell. (***A***) In growth map models, if growth is linear, whereby both winding angle and spiral radius increase linearly in time, the resulting shape is a generalized helix with arithmetic (Archimedean) spiral coiling (Rice 1998), visible here in apical view. The simulation model of Urdy *et al*. (2010) is less amenable to such analysis. But their procedure for constant growth is reminiscent of the recipe for constructing a spiral of Theodorus (Hammer 2016), which tends, again, toward an arithmetic spiral as number of growth steps increases. (***B***) Helicospirals around a shell can be parametrized by position on the perimeter of the aperture, *φ*; azimuth angle, if applicable, or a general parameterization, such as *φ* running from 0 to 1 along the perimeter, or *φ* ∈ [0, *L*] (Goriely 2017), where *L* is the length of the perimeter of the generating curve. This is demonstrated here by adopting the pretty illustrations of *Ecphora* by John S. Spurbeck (Raup 1961). Apical semiangle (left), and helical slope or lead angle (right) vary among spiral paths. *γ* estimated at 0.152 (slightly higher than Raup’s reported value). Measured angles are *β* = 54.45^◦^, 44.75^◦^, 34.71^◦^, 23.02^◦^ and *λ* = 6.65^◦^, 9.17^◦^, 15.16^◦^, 21.64^◦^. Close to theoretical *λ*-values (Eq.[4]) of 6.22^◦^, 8.77^◦^, 12.56^◦^, 20.47^◦^ respectively. (***C***) A similar demonstration using the *Tritia* shell from Johnson *et al*. (2019). *γ* = 0.0825. Upper (suture) spiral: *β* = 30.61^◦^, *λ* = 9.7^◦^. Lower (abapical) spiral: *β* = 9.1^◦^, *λ* = 30.14^◦^. Again, *λ*-values close to theoretical values of 8^◦^ and 29.51^◦^ respectively, as the conical logspiral model predicts.(`R` scripts for calculating the values of angles in panels B and C are available at https://doi.org/10.5281/zenodo.21842256, along with the `svg` image files that contain sampled paths. The original images cannot be posted as part of the CC0-licensed public-domain data repository.)

Taking this multispiral standpoint (also called ‘multivector’; Bayer 1978; McGhee 1978, 1999; Savazzi 1990), helicospiral paths on the shell, designated by position on the aperture’s perimeter, *φ* (Fig. 5B), differ in slope, though expanding at a common rate. A shell’s aperture, therefore, dictates a gradient in lead angle (Fig. 5B,C),

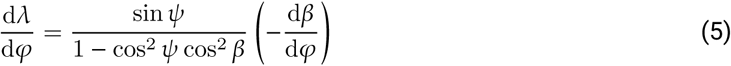

(by applying d d*φ* on Eq.[4]). (Given usual low expansion rates for gastropods, *γ* ≤ 0.2, a simpler approximate expression can be used, d*λ* d*φ* ≈ *γ*/(sin^2^ *β* + *γ*^2^) − d*β* d*φ*; SI.)

This gradient determines how neighboring helicospiral paths on the shell’s surface diverge in lead angle. It is strong close to the coiling axis, at the inner (or columellar) lip (first RHS factor in Eq.[5] goes to csc *ψ* as *β* → 0, and *ψ* is small for gastropods). As apical semiangle increases around the aperture (Fig. 5B,C), i.e., relatively closer to planispiral coiling, the gradient flattens (first RHS factor approaching sin *ψ* as *β* → *π*/2). Consequently, helical slopes are much more sensitive to small changes in *β* on the inner lip of the aperture, i.e., close to the coiling axis, than on its outer margins, as well as overall in more slowly expanding shells (the difference [csc *ψ* − sin *ψ*] grows quickly as *ψ* decreases, at small *ψ*-values).

The gradient in lead angle around the margins of the aperture (Eq.[5]) is a direct geometric consequence of conihelical coiling. That rates and directions of cellular growth, elongation, and division are correlated with this gradient (Johnson *et al*. 2019; Fig. 5C) cannot therefore be taken, on their own, as evidence that such cellular processes are the drivers of coiling. In developmental plasticity parlance, whether the cellular processes are ‘leaders’ or ‘followers’ (West-Eberhard 2003) remains to be determined.

In that context, two hypotheses for the morphogenetic basis of coiling have recently been put forward by Ohta *et al*. (2025). The first, ‘directional cell division’ hypothesis, is an extension of Huxley’s ([1932] 1993) explanation of “growth-centres and growth-gradients in accretionary growth”, backed by modern observations and experimental manipulations (Johnson *et al*. 2019; Ohta *et al*. 2025), and the growth-map school of theoretical morphology (Rice 1998; Hammer & Bucher 2005; Urdy *et al*. 2010; Noshita *et al*. 2016).

The second hypothesis, which Ohta *et al*. (2025) call the ‘mechanical force hypothesis’, states that coiling, and other aspects of shell form, are mostly determined by the interplay of hydrostatic and mechanical forces operating on and within the soft-body, particularly the mantle. This hypothesis relates to Savazzi’s (1990) ‘pneu’, Morita’s (1991; 2003) ‘double membrane tube’ and ‘head-foot guidance mechanism’, and Chirat *et al*.’s (2021) ‘elastic rods’ (also, Goriely 2017). Fluid pressure, action of the columellar muscle, and growth-related accumulation of mechanical stresses in the soft-body tend to press the mantle skirt against particular inner and outer surfaces of the rigid shell, thus guiding future shell accretion.

As pointed out by Vermeij (2002; also Vermeij & Sigwart 2026), unless one subscribes to a “narrow interpretation of the hypothetico-deductive method”, explanations may nevertheless be complementary. Here, I further note that the different morphogenetic hypotheses are often studied at different developmental stages, i.e., during embryonic and/or larval development vs. growth at the juvenile and adult stages (but see Shimizu *et al*. 2013). So also in that respect, the two hypotheses, or sets of explanations, are not mutually exclusive. As discussed by Hutchinson (1989), the initial whorls of the protoconch provide a “nucleating effect” that helps guide the growth of subsequent whorls.

Similarly, van Osselaer & Grosjean (2000) demonstrated, by piecewise fitting of conispirals, significant changes in coiling behavior at the protoconch-teleoconch transition. Cox (1960) and Savazzi (1990) discussed deviated protoconchs, and the significance of protoconch-teleoconch morphological differences has long been recognized by malacologists and palaeontologists (Frýda & Ferrová 2011; Frýda 2012; and references therein). Finally, as Fig. 2 shows for the data of Newkirk & Doyle (1975), embryos and adults differ in basic coiling parameters. Such observations suggest that strong ontogenetic allometry is often confined to the protoconch-teleoconch transition, and that this morphological allometry is coupled to ontogenetic allometry of driving processes.

Finally, the gradient in helical slopes around the aperture, described by Eq.[5], may also assist in understanding the morphology and orientation of columellar folds (Vermeij 2024, 2025). I do not suggest that this gradient in lead angle fully explains the phenomenon. But, as in the case of aperture inclinations (previous section; Fig. 3 and 4), Eqs.[4] and [5] may provide null expectations, to which the observed orientations of columellar folds at different positions on the inner lip can be compared.

### Conclusions

In a humbling example of how, in the natural sciences, even the strongest convictions of the day may turn out to be the outdated dogmas of tomorrow, Moseley (1838), following his seminal analysis of logarithmic spiral shells, concluded that “Providence has subjected the instinct which shapes out each, to a rigid *uniformity* of operation”. Perhaps, once again, D’Arcy Thompson, in his *magnum opus*, was closer to the mark: “The mathematician knows better than we [zoologists, morphologists, biologists] the value of an approximate result” (my brackets); echoing earlier words by Blake (1878) on conihelical geometry serving “as a check on the separate measurements” (see SI).

Here, I explored this premise by studying rarely considered morphological aspects of conihelical geometry, and by demonstrating its usefulness (*i*) in integrating data, obtained by different methodologies (Fig. 2); (*ii*) in identifying natural axes of variation and redundant parameters; (*iii*) in circumventing a pesky long-standing methodological problem that has plagued even recent studies that apply modern methods; (*iv*) to understanding the different types of ontogenetic allometry that break conihelical self-similarity; and (*v*) in assessing adaptational, morphogenetic, and historical interpretations of covariation in coiling parameters. Conihelical geometry explains the observed tight correlation between apical semiangle and expansion rate (Eq.[4]; Fig. 2), regardless of considerations of tight-coiling and whorl-overlap, through plasticity in growth. Replacing the apical semiangle with lead angle, as the relevant morphospace axis, can better help in distinguishing among taxonomic groups, and in understanding and interpreting their morphological differences (Fig. 2B). Additionally, conihelical geometry prescribes a baseline direction for the mantle, columellar muscle, and visceral mass of a shelled gastropod (though there are exceptions; Frýda 2012), serving as reference for other aspects of morphology (e.g., aperture inclinations, columellar folds, growth gradients).

Despite such a constraint, the head-foot mass and opening of the shell are grown and maintained parallel to the substrate. One can call that homeostasis. One can call that adaptation — for defense, postural stability, aestivation, etc. One can consider that an innovation that lead to the diversity explosion in conchiferans (Vermeij & Sigwart 2026). One can see it as a constructional consequence of shell and substrate together directing the soft body, its development, and subsequent shell accretion. Physiology and behavior translate morphology into ecology (Ricklefs & Miles 1994). Equally, through development, ecology and life habit feed back into morphology (e.g., Thompson 1999). In natural history, “explanations tend to be complementary, not exclusive”, and adopting a “narrow interpretation of the hypothetico-deductive method” may be counterproductive (Vermeij 2002). To quote from another *magnum opus*, “the role of the environment as an agent of development, not just selection, in the evolution of all forms of life.” (West-Eberhard 2003). Also this principle is, at times, “apt to be forgotten” (see a previous section).

A recent astronomy paper commented that “It is remarkable that after over 170 years of observations of spiral arms in galaxies our understanding of them remains incomplete.” (Masters *et al*. 2019). The first sighting of galactic spiral arms, by Lord Rosse in 1846, coincided with the first incarnation of the recurring debate over the “true spiral” (Naumann 1845; Raup 1966) of shell-building organisms. Another aspect of not getting bogged down in too narrow application of hypotheses-testing is admitting that organic structures seldom exhibit “Platonic ideals” (McGhee 2001), but that such ideal models may still hold well, as useful first approximations (Aldridge 1998; van Osselaer & Grosjean 2000), and “by predominant relative frequency” (Gould 2002). As I demonstrated here, the parameters and geometric relationships that conihelical geometry prescribes are still very much relevant for interpreting ontogenetic and intra– and interspecific (co)variation, including allometry.

The gastropod shell is a permanent record of an animal’s ontogeny, while spiral arms of galaxies may be ephemeral patterns, constantly forming and dissipating, albeit on a vastly different time scale. Yet, the variability and many irregularities in galactic spiral arms have not prevented astronomers from applying the pitch angle (here, expansion angle) as a useful parameter, in classifying galaxies, in studying relations to other variables (e.g., mass), and in testing theories of formation and evolution of such spirals. Perhaps natural history and evolutionary biology can borrow a page from a fellow historical science.

## Acknowledgements

Images in Fig. 5B,C from Raup (1961) and Johnson *et al*. (2019), reproduced here under fair use and conditions of the PNAS license. The original illustrations of *Ecphora* (Fig. 5B) are by John S. Spurbeck.

## DATA AND SOFTWARE AVAILABILITY

All data, code, and images are available at https://doi.org/10.5281/zenodo.21842256. Web application for theoretical morphology of coiled shells (Fig. 1A,B) is available at https://doi.org/10.5281/zenodo.22165306.

