## Supplementary information PDF for "Galactic arms, and devil’s toenails: a synthesis of coiled morphology, applied to gastropods"

### Supplementary information for "Galactic arms, and devil's toenails: a synthesis of coiled morphology, applied to gastropods"

Ido Filin [1*]

[1] Independent researcher.

https://orcid.org/0000-0001-6231-0029

### Contents

**S1 General planispiral coiling** 2

**S2 Estimating planispiral and helicospiral arclength** 2

**S3 Approximation errors** 4

**S4 Measurement errors, coiling axes misalignment, regression weighting, and specimen repeatabilities** 4

**S5 Data analyses** 9

**S6 Model selection procedures** 12

**S7 Optimal control of planispiral growth** 14

**S8 Web application** 16

**S9 Quote from Blake 1878** 16

**S10 Supplementary references** 17

#### S1 General planispiral coiling

Burnaby (1965, 1966) previously defined an allometric planispiral through a power-law relationship of arclength to radius, (using the notation of this study) $\ell = \ell(r) = c_0 + c_1 r^k$, $c_0$, $c_1$, and $k$ constants. Much of the early *Gryphaea* debate revolved around the value of $k$, in different species and strata. For logarithmic planispirals, $c_0 = 0$, $k = 1$, and $c_1 = \sec\alpha = \csc\psi$; the familiar $\ell = r/\cos\alpha$ of logspirals.

For any spiral plane curve, the increment in winding angle is related to increment in arclength through $\mathrm{d}\theta = \mathrm{d}\ell\cos\psi/r$. Coiling rate, or coiling tightness, $\mathrm{d}\theta/\mathrm{d}\ell$, is $\cos\psi/r$. If both $\psi$ and $r$ depend on $\ell$, net change in winding angle along a section of the spiral will be given by the integral $\theta_{12} = \int_{\ell_1}^{\ell_2} \frac{\cos\psi(\ell)}{r(\ell)}\,\mathrm{d}\ell$. Because $\mathrm{d}\ell/\mathrm{d}r = \csc\psi$, the integrand becomes $\cot\psi\,\mathrm{d}r\,/r$. For fixed $\psi$, this just integrates to $\theta_{12} = \tan\alpha\ln(r_2/r_1)$, the equiangular logarithmic spiral. In other cases, integrating by parts gives

$$\theta_{12} = \left(\cot\psi\ln r\right)\Big|_{r_1}^{r_2} + \int_{r_1}^{r_2} \psi'\csc^2\psi\ln r\,\mathrm{d}r \tag{S1}$$

($\psi' = \mathrm{d}\psi/\mathrm{d}r$), which again shows that the equiangular case, $\psi' = 0$, results in the logarithmic spiral expression, $\theta_{12} = \cot\psi\ln(r_2/r_1)$. The second RHS term can be rewritten as $\int_{\ell_1}^{\ell_2}(\mathrm{d}\psi/\mathrm{d}\ell)\ln r(\ell)\,\mathrm{d}\ell$, and again through integration by parts,

$$\theta_{12} = \left(\cot\psi\ln r\right)\Big|_{r_1}^{r_2} + \left(\psi\ln r\right)\Big|_{r_1}^{r_2} - \int_{r_1}^{r_2}\frac{\psi}{r}\,\mathrm{d}r\,. \tag{S2}$$

By defining, for the last RHS term, the $\ln r$-weighted mean expansion angle along the spiral section,

$$\langle\psi\rangle_{12} = \frac{\int_{r_1}^{r_2}\frac{\psi}{r}\,\mathrm{d}r}{\int_{r_1}^{r_2}\frac{1}{r}\,\mathrm{d}r} = \frac{\int_{\ln r_1}^{\ln r_2}\psi\,\mathrm{d}(\ln r)}{\ln r_2 - \ln r_1}, \tag{S3}$$

Eq.[2] of the main text is finally obtained.

In the case of expansion angle increasing or decreasing linearly with $\ln r$, Eq.[1] of the main text can be integrated into a closed form, $\theta = \int_0^{\ln(r/r_0)}\cot(a+bx)dx = \frac{\ln\sin(a+b\ln(r/r_0))}{b} - \frac{\ln\sin a}{b}$. Further trivial derivations result in Eq.[3]. Taking the Taylor series of Eq.[3] around $a = 0$, to the first non-zero order in $a$ (linear term), one obtains

$$\ln r = \ln r_0 + \frac{a}{b}\left(e^{b\theta} - 1\right) + O\left(a^3\right), \tag{S4}$$

which can also be used as a regression model, if expansion angle remains small throughout the relevant section of the planispiral.

#### S2 Estimating planispiral and helicospiral arclength

In this section I derive expressions for estimating arclength from a sequence of successive radii separated by half-revolutions, used in subsequent data analyses. If the diameter of a planispiral segment, spanning a half revolution, is known, i.e. $r_1 + r_2$ given and $r_1$ and $r_2$ separated by $\theta_{12} = \pi$, the arclength of a logspiral segment of constant expansion angle, $\psi = \mathrm{const}$, is given by $(r_1 + r_2)\tanh(\pi\gamma/2)\csc\psi$, where $\gamma = \tan\psi$. The factor multiplying $(r_1 + r_2)$ can be rewritten as $\left(\sqrt{1+\gamma^2}/\gamma\right)\tanh(\pi\gamma/2)$, which is a decreasing monotonic function in $\gamma \geq 0$, with a maximum value of $\pi/2$ at $\gamma = 0$.

Assuming that this planispiral segment, with arclength $\ell_{12}$, can be considered logspiral, i.e., $\psi$ and $\gamma$ constants over a half revolution, and limiting values of expansion rate to the values most commonly observed in gastropods, $\gamma \in [0, 0.2]$, I obtain the condition

$$1.5512(r_1 + r_2) \leq \ell_{12} \leq \frac{\pi}{2}(r_1 + r_2) \qquad \text{(S5)}$$

($\pi/2 =$1.5708). Taking the approximation

$$\ell_{12} \approx \frac{\pi}{2}(r_1 + r_2) \qquad \text{(S6)}$$

therefore results in relative error of no more than 1.27%, which is negligible compared to other error components in the data analyses, discussed below. Based on optimal control paths (Optimal control section, below), composed of circular paths, line segments, and logspiral segments, a higher range of relative error in arclength is obtained, 7.5%. However, that is still a relatively small contribution, well within the range of intra-group differences in estimated lead angle.

Given a sequence of successive centroids, $r_1, r_2, r_3, \ldots, r_n$, separated by half revolutions, $\theta_{i,i+1} = \pi$, the total planispiral arclength between $r_1$ and $r_n$ is approximated by

$$\ell_{1n} \approx \frac{\pi}{2}\left(r_1 + r_n + 2\sum_{j=2}^{n-1} r_j\right). \qquad \text{(S7)}$$

Again, with an error of no more than 1.27%.

For completeness, the approximation error can also be estimated from the Taylor series of $(\sqrt{1+\gamma^2}/\gamma)\tanh(\pi\gamma/2)$, which, truncating to second-order, is given by $\frac{\pi}{2} + \gamma^2\left(\frac{\pi}{4} - \frac{\pi^3}{24}\right) + O(\gamma^4)$. The first significant order is the constant $\frac{\pi}{2}$. The second-order term, therefore, provides an estimate of the error when taking the approximation in Eq.[S6], which is 0.02026 for $\gamma = 0.2$, or 1.29%.

The average helical slope along a helicospiral segment can then be estimated from centroids of sagittal cross-sections of shells by dividing the longitudinal difference, i.e., lead, $z_n - z_1$, by arclength, $\ell_{1n}$,

$$\overline{\tan\lambda} \approx \frac{2}{\pi}\frac{z_n - z_1}{r_1 + r_n + 2\sum_{j=2}^{n-1} r_j}. \qquad \text{(S8)}$$

For two successive centroids, $(r_1, z_1)$ and $(r_2, z_2)$ this approximate relation is identical to the formula for the lead angle of a screw where the numerator, $z_2 - z_1$ is the screw's lead, and the denominator, $r_1 + r_2$ is the diameter of the screw's helical thread.

Three-dimensional helicospiral arclength can be estimated, in a similar fashion to Eq.[S7], by summing the increments of conihelical segments. Denoting such three-dimensional arclength by $s$, the estimate for arclength of a helicospiral segment, $s_{1n}$, is given by

$$s_{1n} = \sum_{j=2}^{n}\sqrt{\left(\ell_{1j} - \ell_{1,j-1}\right)^2 + \left(z_{1j} - z_{1,j-1}\right)^2}, \qquad \text{(S9)}$$

or, for slow expansion, $\gamma \leq$ 0.2,

$$s_{1n} \approx \sum_{j=2}^{n}\sqrt{\frac{\pi^2}{4}\left(r_j + r_{j-1}\right)^2 + \left(z_{1j} - z_{1,j-1}\right)^2}. \qquad \text{(S10)}$$

Such empirically estimated three-dimensional arclength was used, in this study, only in model selection of allometry models for aperture size, i.e., $A(s)$ (see Model selection procedures below).

#### S3 Approximation errors

Approximation errors for arclength and for the nonlinear planispiral linear-log-$\psi$ regression formula (Eq.[3] in main text) are discussed above, in their relevant sections (see Eqs.[S4] and [S7]).

For small $\gamma$, an approximate form of Eq.[4] (main text) is given by $\tan\lambda\tan\beta \approx \gamma$, and an approximate form of Eq.[5] (for the gradient in lead angle around the generating curve) is given by

$$\frac{\mathrm{d}\lambda}{\mathrm{d}\varphi} \approx \frac{\gamma}{\sin^2\beta + \gamma^2}\left(-\frac{\mathrm{d}\beta}{\mathrm{d}\varphi}\right). \tag{S11}$$

It is easy to show that, in both cases, the ratio of the approximate expression to the exact expression is $\sqrt{1+\gamma^2}$ (or $\sec\psi = 1/\cos\psi$), which is a monotonically increasing function of $\gamma$. Consequently, in the domain $\gamma \leq 0.2$, the relative error is no more than 2% ($\sqrt{1+0.2^2} = 1.0198$).

#### S4 Measurement errors, coiling axes misalignment, regression weighting, and specimen repeatabilities

Fig. S1 presents several examples of specimens, of different species and numbers of sampled cross-sections/centroids, from the Collins *et al.* (2021a) dataset. Additionally, the figure presents the estimated conihelical centerlines, as estimated in this study. Fig. S2 shows several images of the resampled *Tylospira coronata* specimen `697a`, and resampled *Pelicaria vermis* specimen `060`. Within-specimen uncertainty in $\tan\psi$- and $\tan\lambda$-estimates are then estimated, pointing at small effects. OLS-estimated $\gamma$ ranged from 0.083 to 0.092 for `697a` (10 samples), and from 0.0802 to 0.0823 for `060` (9 samples). Mixed-effect modeling with `lmer` (Bates *et al.* 2015), applying sample-dependent random intercepts, estimated the standard deviation associated with (re)sample at 0.0014 for `697a` (predicted $\gamma$-values from 0.0874 to 0.090), and too small to be estimated (i.e., singular model) for `060`. Overall, effect of per-specimen digitization and measurement error is clearly quite small, about 2%. Similar results are obtained for $\tan\lambda$-estimates (from 0.27 to 0.298 for `697a`, and from 0.343 to 0.3475 for `060`).

Realigning coiling axes according to the centroids bisector (e.g., Davoli & Russo 1974) produced, in the vast majority of cases, barely perceptible corrections (Fig. S3). In addition, I tried different weighting schemes for per-specimen centroids, in $r(\theta)$-, $y(\ell)$- and $y(r)$-regressions, to check for robustness of results, and to further assess estimation errors of coiling parameters (either no weighting, or weights by aperture area, $A$). Differences in expansion- and lead-angle estimates by different weighting were the strongest component of estimation error, in comparison to effects of resampling and coiling axis misalignment, but still produced little overall effect (see also Fig. S4 and S5). Specimen repeatabilities were estimated with `rptR` (Stoffel *et al.* 2017) to be very high, both within groups and in total, as reported in main text.

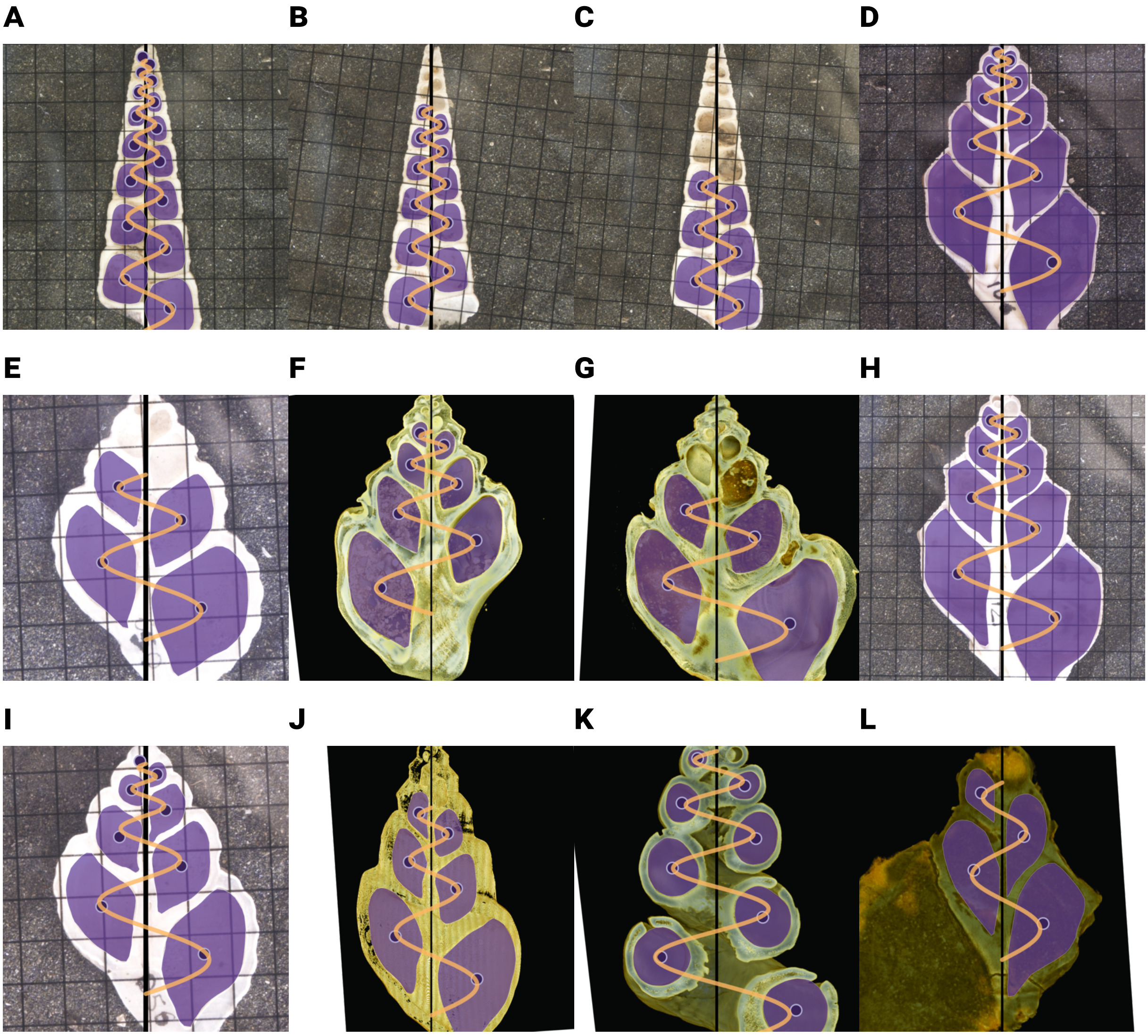

Figure S1 : Some examples of sectioned shells from the dataset of Collins *et al.* (2021*a*), and estimated conihelical centerline paths of the present study. As part of the reanalysis done in this paper, I generated `svg` images, with highlighted apertures (shaded purple), and aperture centroids (dark purple dots), coiling axes (solid black line), and estimated conihelical centerline (light orange). Compared to the originals, the generated `svg` images have been zoomed in on the specimens and rotated, such that the measured coiling axis (black line) is vertical. (***A,B,C***) Specimens of *Maoricolpus roseus* (Turritellidae): specimen IDs `071`, `082` and `073` respectively. (***D,H***) Specimens of *Struthiolaria papulosa* (Struthiolariidae): specimen IDs `106` and `121` respectively. (***E,I***) Specimens of *Pelicaria vermis* (Struthiolariidae): specimen IDs `050` and `056` respectively. (***F,G***) Specimens of *Tylospira coronata* (Struthiolariidae): `P135697e` and `P135835g` respectively. (***J***) Specimen of *Tylospira scutulata* (Struthiolariidae), `VM989`. (***K***) Specimen of *Tenagodus anguinus* (Siliquariidae), `P325996-4`. (***L***) Specimen of *Struthioptera camachoi* (Aporrhaidae), `WM15706`.

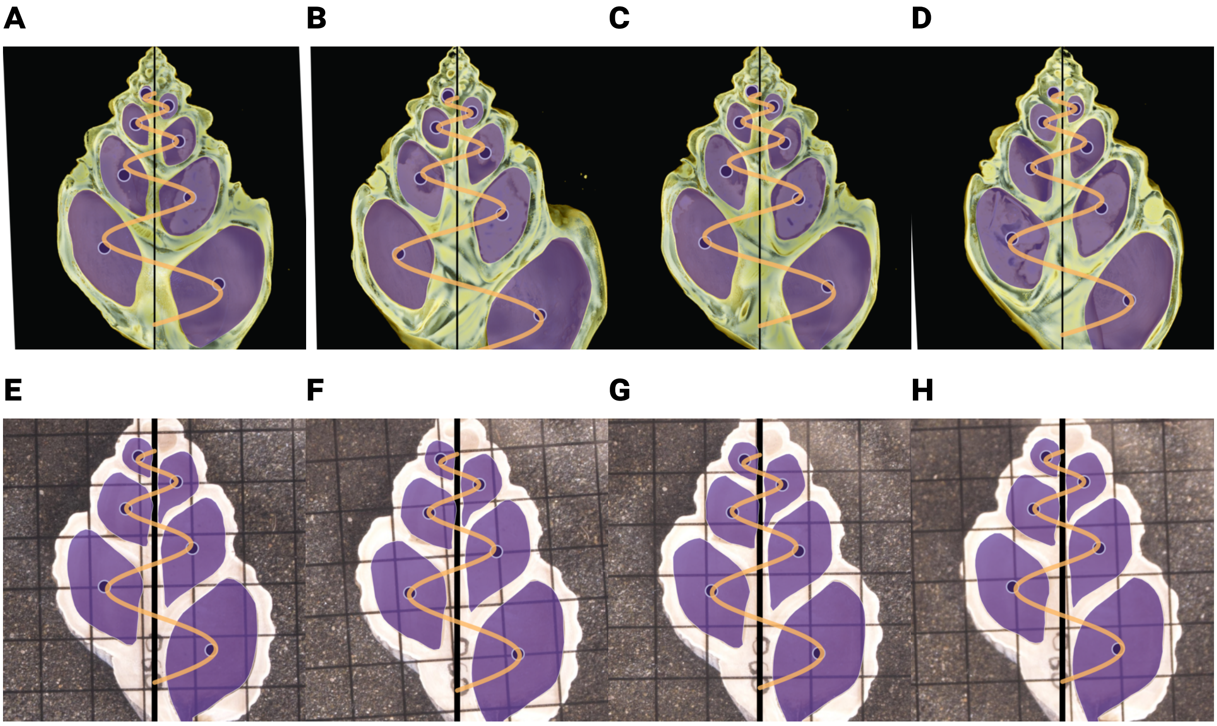

S132 Figure S2 : (***A–D***) Four of the ten resampling replications of the *Tylospira coronata* specimen `697a`. (***E–H***) Four of
S133 the nine resamples of the *Pelicaria vermis* specimen `060`. These specimens were resampled by Collins *et al.* (2021*b*),
S134 for the purpose of estimating digitization error. They can be used to assess magnitude of effects of such sampling
S135 error on coiling parameter estimates.

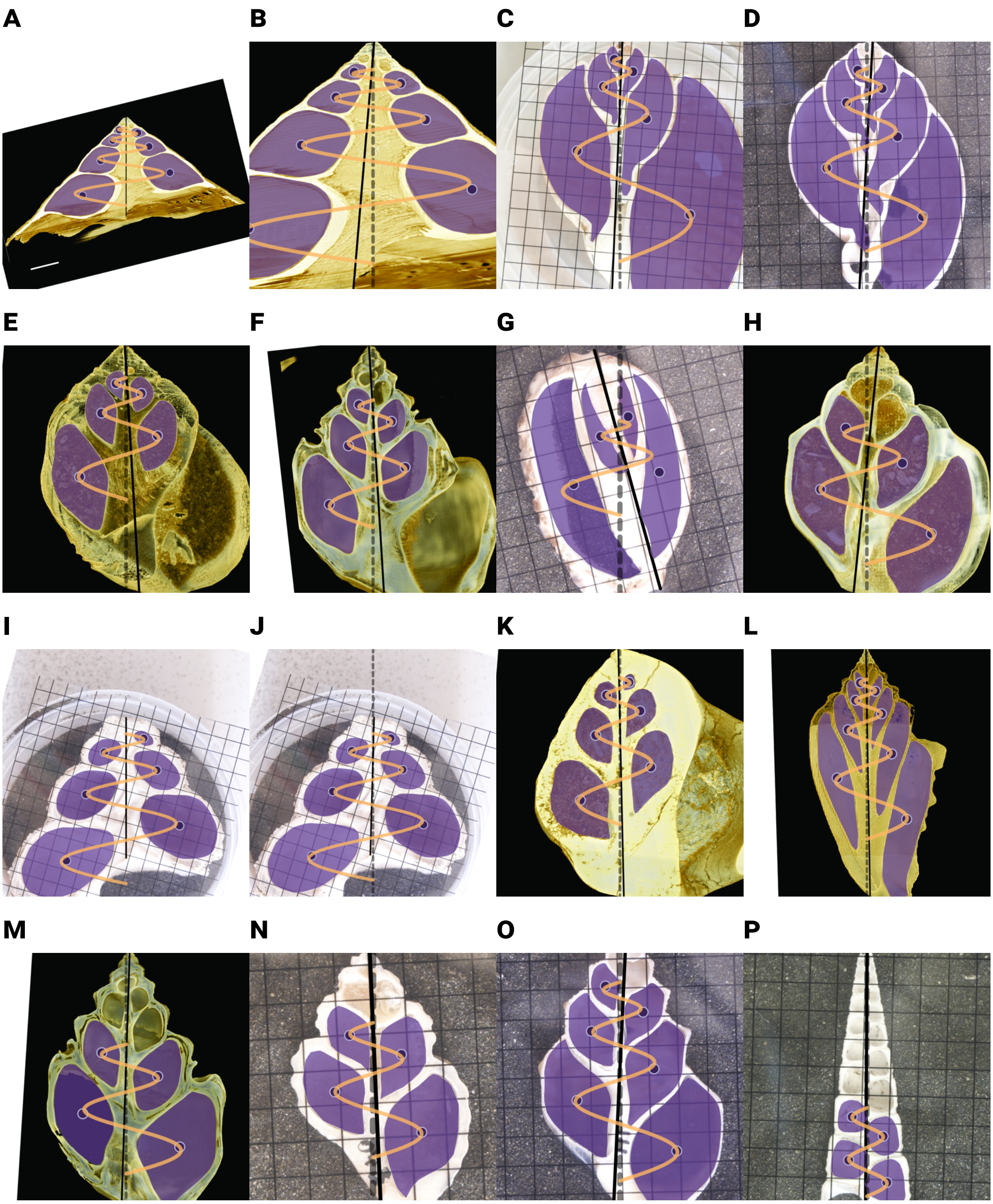

Figure S3 : Some images exhibit clear visual signatures of mismeasured coiling axes. The clearest example of that is (***A***) the *Xenophora exuta* specimen `VM990`. Axis misalignment is reflected in centroids on one side being systematically closer to the coiling axis, compared to those on the opposite side. (***B***) Applying the bisector method (Davoli & Russo 1974) to realign such mismeasured axes clearly corrects the effect. Significant bisector-based corrections were observed in few other specimens: (***C***) *Tonna variegata* `Tvariegata`; (***D***) *Semicassis pyrum* `104_1`; (***E***) *Perissodonta nordenskjoldi* `WM12400`; and (***F***) *Tylospira coronata* `P135835f`. In some cases the bisector algorithm failed to correct the coiling axis, by producing and axis that clearly does not pass through the apex of the shell. This effect was confined to some specimens with only 4 centroid points, such as (***G***) the *Cypraea* specimen `95`, and (***H***) *Tylospira gilli* `P134825`, or to the open-coiled *Tenagodus anguinus*. In the vast majority of specimens however (131 out of 164) the recalculated axis was barely distinguishable visually from the original manually measured axis. For example, (***I,J***) *Cookia sulcata* `Csulcata`; (***K***) *Conchothyra parasitica* `AG6377`; (***L***) *Strombus pipus* `VM988`; (***M***) *Tylospira coronata* `P135835k`; (***N***) *Pelicaria vermis* `048`; (***O***) *Struthiolaria papulosa* `111`; and (***P***) *Maoricolpus roseus* `072`.

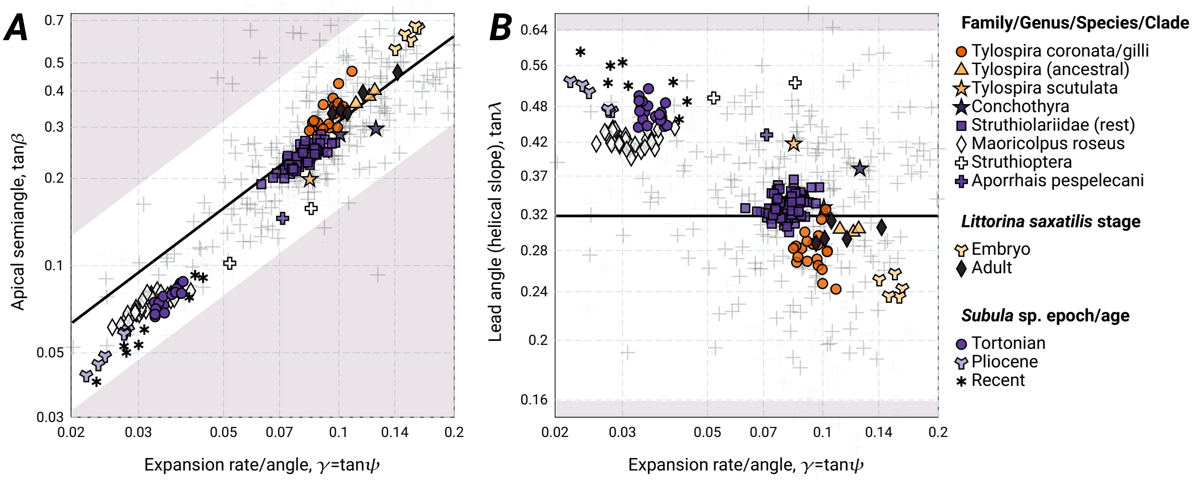

S148 Figure S4 : A version of Fig. 2 of main text, after realigned coiling axes.

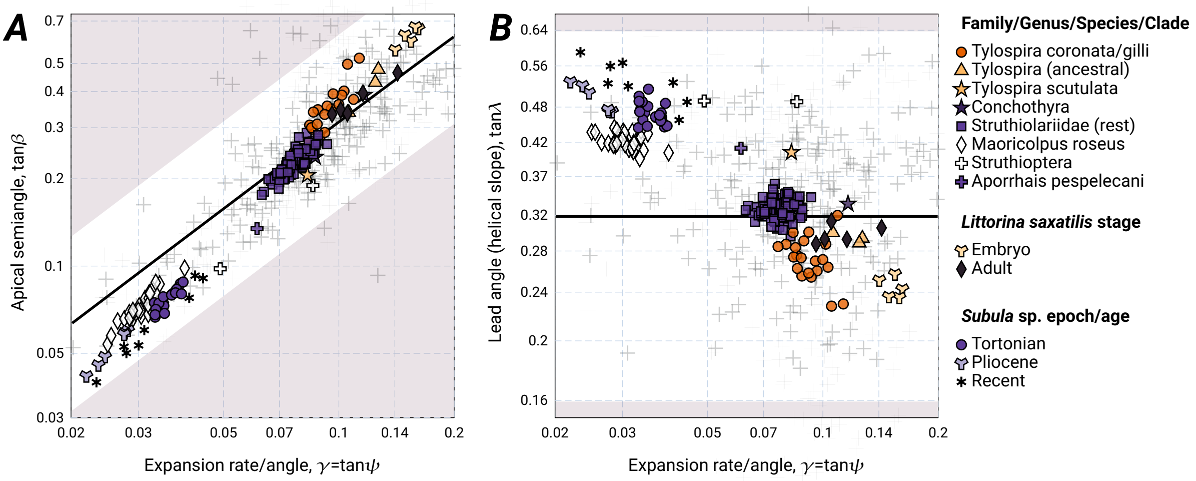

S149 Figure S5 : A version of Fig. 2 of main text, after regression weighting according to aperture areas.

#### S5 Data analyses

##### *S5.1 Coiling parameters for the Collins* et al. *dataset*

I obtained the data of Collins *et al.* (2021*b*) from the data repository of that study (https://doi.org/10.5061/dryad.p5hqbzknw). For convenience in subsequent analyses, as well as visual inspection and verification of fitted conihelical curves, I wrote an `awk` script that scans the original `tps` file of raw morphometric measurements, renames the original `jpg` images, calls `R` for some statistical analyses and potentially coiling axis realignment, collects and merges images' metadata and per-specimen and per-aperture data into new `csv`-files, and draws per-specimen `svg` images from the original `jpg` files, data, and estimated conihelical centerline. More details are available inside code at https://doi.org/10.5281/zenodo.21842256. All (renamed) images, `csv` and `svg` files, and code (e.g., `awk`- and `R`-scripts) are available at that repository.

Each specimen (or specimen resample), in the Collins *et al.* dataset, contains a sequence of aperture centroids and areas, $(r_i, y_i, A_i)$, at half-whorl intervals, i.e., $\theta_{i-1,i} = \pi$. From these, planispiral arclengths, $\ell_i$, and total (three-dimensional) arclengths, $s_i$, are obtained from Eqs.[S7] and [S10].

OLS linear and quadratic regressions were conducted with `R`'s basic `lm` function. Nonlinear models were fitted with `nlsLM` from the `minpack.lm` package (Elzhov *et al.* 2023). Model II linear regressions were conducted either with eigenanalysis of covariance matrices (using `cov.wt`), with the `lmodel2` package (Legendre 2024), or with the `smatr` package (Warton *et al.* 2012).

For expansion rate, $\gamma$, estimates from either linear regression or Model II major-axis (MA) regression produced practically identical values, as reported in main text. $R^2$-values again were very high, as discussed in main text. Out of the total 164 specimens, only six exhibited marked deviation from a logspiral in $\ln r$-$\theta$ regression, as measured by $R^2_{adj}$ lower than 0.9. Those six specimens either belonged to the open-coiled *Tenagodus anguinus* or had only 4 centroid points (e.g., Fig. S3N,P). Only the *Tenagodus* specimens had $R^2_{adj}$ lower than 0.84.

Per-specimen lead angles were estimated from $y(\ell)$-regressions, either OLS or Model II MA. Values of $R^2$ and $R^2_{adj}$ were again very high, as reported in main text. Of the three main clades (148 of 164 specimens) no specimen had $R^2_{adj}$ below 0.97.

Results of quadratic and nonlinear regressions are discussed in detail in the main text. In this supplemental document, I only present Fig. S6, as a visual aid to assess the magnitude of this quadratic/nonlinear effects, which are quite weak.

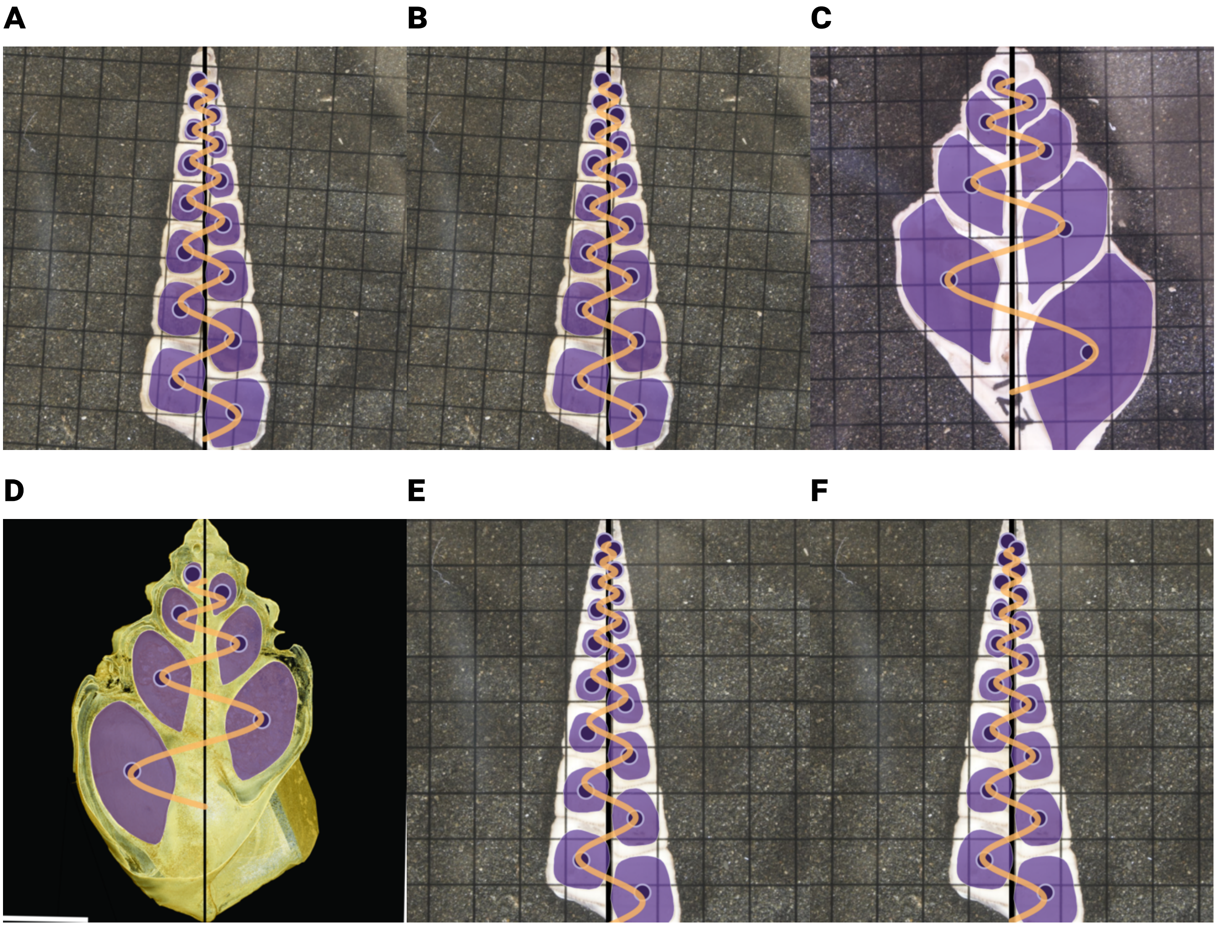

Figure S6 : Examples of allometric centerline coiling, identified after Bonferroni-Holm correction. (***A***) Specimen `079` was the only specimen having trends in both lead angle and expansion angle. The trend in lead angle can be appreciated by comparing to (***B***) the centerline estimated with aperture-size-based weighted regression. Although statistically significant, the effect is quite weak. (***C***) For specimen `127` (*S. papulosa*), expansion decreases ontogenetically, as evident from the centroid position relative to estimated conihelical curve. Still, the effect is again small. (***D***) Ontogenetic change in lead angle is evident from centroid positions with respect to estimated curve. For this image of the *T. coronata* specimen `P135697d`, conihelical centerline was estimated with aperture-size-based weighted regression, to make the change in lead angle at younger (smaller) whorls more clear. (***E,F***) *M. roseus* specimen `100`, with no weighting (E), and aperture-size-weighting (F), demonstrating again the slight ontogenetic change in lead angle at younger whorls.

##### *S5.2 Mixed-effect growth models*

To facilitate interspecific comparisons, I fitted a mixed-effect linear growth model of $\ln A$ against $\ln r$, with random per-specimen intercept and slope, and the three clades of *Maoricolpus*, *Tylopspira+Perissodonta* and rest of the Sturthiolariidae as fixed effects (Grimm *et al.* 2017). Data for this mixed model was thus limited to specimens of *Maoricolpus* and Struthiolariidae — 148 out of the total 164 specimens. Analysis was conducted `lmer` of the R-package `lme4` (Bates *et al.* 2015), as well as with `nlme` of the `nlme` package (Pinheiro *et al.* 2025). Estimates and confidence intervals for aperture allometry, reported in the main text, are from `nlme`. Estimates and confidence intervals from `lmer` were very similar and did not change the qualitative conclusions in the main text.

Similarly, as reported in the caption of Fig. 2 in the main text, linear mixed-effect growth models for

$y(\ell)$ provided clade-specific estimates and confidence intervals of $\tan\lambda$. I additionally examined quadratic terms in the mixed models (with `lmer`). Although quadratic fixed effects were statistically significant (by applying `anova`), inspection of predicted values and residuals showed practically no improvement by these quadratic term. This conclusion is further corroborated by examining $R^2$-values, using the `r2` function of the `performance` package (Lüdecke *et al.* 2021).

##### *S5.3 Analysis of aperture inclinations in the Noshita et al. dataset*

Limiting data to the main sequence of coiling parameters, $\tan\psi \leq 0.2$ and $1/2\pi \leq \tan\lambda \leq 2/\pi$, one-sample $t$- and Wilcoxon tests, separately on the three habitat groups and on (relative) outward, $\Gamma^*$, or downward, $\Delta^*$, inclinations revealed that, for land snails, mean and median inclinations are both positive and significantly different than zero ($\langle\Gamma^*\rangle = 0.2687$; $\langle\Delta^*\rangle = 0.2183$; $p < 0.0001$). For freshwater species, only values for relative downward angle, $\Delta^*$, were significantly different than zero, and even then practically nil (mean -0.053, $p = 0.011$; median -0.06 $p = 0.0032$). For marine species, only values for $\Gamma^*$ were significant (mean 0.2242; median 0.2154; $p < 0.0001$); while median (but not mean) $\Delta^*$ was also significantly different from zero ($p = 0.024$), its value was very small, -0.054.

Contingency table analyses on the signs of $\Gamma^*$ and $\Delta^*$ ($\chi^2$-test), separately for each habitat group, revealed the differences in excesses of distribution among the different habitats, as presented in the main text, and were fully consistent with the results of $t$- and Wilcoxon tests. (For freshwater, $\chi^2 = 12.85$, $p = 0.005$; For land, $\chi^2 = 59$, $p <$ `1e-12`; For marine, $\chi^2 = 79.96$, $p <$ `2.2e-16`.) These conclusions are further corroborated by applying analysis of variance on distance matrices in $(\Gamma^*, \Delta^*)$-morphospace with `adonis2` and `pairwise.adonis2` (Oksanen *et al.* 2025, Martinez Arbizu 2017).

For testing relationships between inclinations (absolute or relative) and expansion and lead angles, I applied linear regression with `lm`, SMA model II regression with the `smatr` package (Warton *et al.* 2012), and linear mixed-effect models (LMMs) with `lme4` and related packages (Bates *et al.* 2015, Kuznetsova *et al.* 2017, Lüdecke *et al.* 2021). Overall, in the freshwater realm, absolute inclinations, $\Gamma$ and $\Delta$, are independent of lead/expansion angles, being distributed close to zero. Marine and land species share the same inverse relationship (i.e., common slopes) between downward inclination, $\Delta$ or $\Delta^*$, and lead angle, $\lambda$, and direct relationship with $\psi$. The latter effect, however, is interpreted as a statistical artifact of the negative correlation in land snails between outward and downward inclination ($r = -0.55$ for $\Gamma$ and $\Delta$; $r = -0.53$ for $\Gamma^*$ and $\Delta^*$), as clearly visible in Fig. 3*A*,*B*. These relationships, however, practically disappeared in the marine realm when accounting for taxonomic family as random intercepts with LMMs.

For outward inclination, $\Gamma$ and $\Gamma^*$, the only effect was a negative relationship with expansion angle, $\psi$, for land snails. This effect also disappeared after accounting for random intercepts by taxonomic family.

Model II SMA regressions similarly revealed that, only for land snails, outward inclination, $\Gamma$ or $\Gamma^*$, decreases with expansion angle, $\psi$. Otherwise, outward inclinations were uncorrelated with expansion or lead angles. Downward inclination, $\Delta$ or $\Delta^*$, decreased with lead angle, $\lambda$, and increased with expansion angle, with common slopes for marine and land species. In the freshwater realm, only a negative $(\Delta^*, \lambda)$ correlation was statistically significant.

As Fig. 4 in the main text shows, the differences in distributions of taxa in $(\Gamma^*, \Delta^*)$-morphospace and the negative relationship between $\Delta^*$ and $\lambda$ are to a large extent effects of taxonomic family. After removal of family effects (purely random-intercept LMMs), the only consistent relationship for within-family residuals is the negative correlation between $\Delta^*$-residuals and $\lambda$-residuals. This is a highly significant relationship with common slope and elevation, shared by all three habitat realms. (Freshwater, $R^2 = 0.1$, $p = 0.001$; Land, $R^2 = 0.44$, $p =$ `1.27e-11`; Marine, $R^2 = 0.27$, $p <$ `2.22e-16`; common slope $-2.8$ $[-2.55, -3.1]$; elevation not significantly different than zero, for obvious reasons).

To test for robustness of patterns several bootstrap procedures were employed. First, for LMMs with `lmer` the built-in bootstrap machinery in `lme4` was used, to obtain confidence intervals for estimated coefficients. For SMA model II regressions of residuals, after removal of taxonomic family effects, resampling of the data, either complete-dataset-wise, or separately within taxonomic families, including removal of small-sample-size families and downsampling large families, to even per-family sample sizes (e.g., reducing family-size range from 1 – 32 to 4 – 10), revealed that the only consistently robust pattern was the negative relationship between $\Delta^*$-residuals and $\lambda$-residuals, common to all three habitats, and potentially a negative relationship between $\Gamma^*$-residuals and $\psi$-residuals only in the marine realm.

#### S6 Model selection procedures

Model selection procedure follows Collins *et al.* (2021*b*) and `R` code, adapted and modified from Collins *et al.* (2021*a*). The procedure fits all models to all specimens, and compares models using Mean $R^2$ and Mean relative likelihoods, where relative likelihood is by comparing each model to the per-specimen best-performing (based on $AIC_c$-values). Means are taken over all 148 specimen of of the three main clades of the dataset. Results are presented in Table S1.

Linear models are fitted with `lm` and nonlinear models with `nlsLM` (Elzhov *et al.* 2023). $AIC_c$-values are obtained with the `AICcmodavg` package (Mazerolle 2023). Linear $y(r)$, $y(\ell)$ and $\ell(r)$ are the preferred models, as expected for conical helices (logarithmic conispirals).

Analysis of base planispirals $r(\theta)$, again revealed the logspiral as the preferred model, with linear-log expansion angle, Eq.[3] of main text, as second. Again, a strong corroboration for the conihelical model, when combined with the evidence for linear $\ell(r)$.

For aperture size, $A$, I examined models of the form $A(\theta)$, $A(r)$ and $A(s)$, where $r$ is centerline radius, and $s$ is empirically measured centerline arc length, from Eq.[S10]. The best performing model is the power-law in radius. The next best is the exponential in $\theta$. These models all point to a general (geometrically sensible) allometric relationship between aperture area and centerline radius, where the isometric case is $A \propto r^2$.

Table S1: Model selection for conical envelopes, centerline spiral radii, planispiral arclength, and aperture area. Formulas of models are presented in a form that is loosely based on R's formula format. Mean $R^2_{adj}$ are given, as well as the mean (per-specimen) relative likelihood of each model, and similarly obtained mean relative likelihoods after removing the best model. The model-dependent symbols $k$, $c_0$ and $c_1$ represent general fitted parameters, not otherwise defined in the main text. Linear models with log-transformed (response or predictor) variables implicitly include a small correction value of $0.0001$, to avoid $\ln(0)$ (as in Collins *et al.* 2021*b*).

| Model | Mean $R^2$ | Mean relative likelihood | Mean relative likelihood (best removed) |
|---|---|---|---|
| **Spiral radius**: $r(\theta)$ | | | |
| Log, $r \sim \ln\theta$ | 0.27731 | <0.00001 | 0.00002 |
| Power-law, $\ln(r) \sim \ln\theta$ | 0.44798 | 0.00006 | 0.05128 |
| Linear, $r \sim \theta$ | 0.95425 | 0.00102 | 0.04779 |
| `nls` Power-law, $r \sim (c_1\theta + c_0)^k$ | 0.97759 | 0.00459 | 0.00494 |
| Quadratic, $r \sim \texttt{poly}(\theta, 2)$ | 0.97806 | 0.00488 | 0.00502 |
| `nls` Logspiral, $r \sim r_0 e^{\gamma\theta}$ | 0.97863 | 0.00196 | 0.25473 |
| Logspiral, $\ln(r) \sim \theta$ | 0.97870 | 0.67020 | – |
| log Quadratic, $\ln(r) \sim \texttt{poly}(\theta, 2)$ | 0.98141 | 0.35415 | 0.61395 |
| `nls` Planispiral, $\ln(r) \sim \ln r_0 + \frac{\arcsin\left(e^{b\theta}\sin a\right) - a}{b}$ | 0.98156 | 0.42178 | 0.69260 |
| **Conical envelope**: $y(r)$ | | | |
| Linear $y(r)$, $y \sim r$ | 0.97610 | 0.77857 | – |
| Quadratic $y(r)$, $y \sim \texttt{poly}(r, 2)$ | 0.97740 | 0.32575 | – |
| **Lead**: $y(\ell)$ | | | |
| Linear $y(\ell)$, $y \sim \ell$ | 0.99798 | 0.76767 | – |
| Quadratic $y(\ell)$, $y \sim \texttt{poly}(\ell, 2)$ | 0.99853 | 0.31446 | – |
| **Planispiral arclength**: $\ell(r)$ | | | |
| Linear, $\ell \sim r$ | 0.97826 | 0.77349 | – |
| Quadratic, $\ell \sim \texttt{poly}(r, 2)$ | 0.97962 | 0.34216 | – |
| **Aperture area**: $A(\theta)$, $A(s)$, or $A(r)$ | | | |
| Power-law, $\ln(A) \sim \ln\theta$ | 0.50073 | 0.00008 | 0.00020 |
| Power-law, arclength, $\ln(A) \sim \ln(s)$ | 0.54636 | 0.00010 | 0.00028 |
| `nls` Power-law, $A \sim (c_1\theta + c_0)^k$ | 0.80889 | 0.04823 | 0.04823 |
| Log, $A \sim \ln\theta$ | 0.84739 | <0.00001 | <0.00001 |
| Linear, $A \sim \theta$ | 0.89288 | <0.00001 | <0.00001 |
| Exponential, arclength, $\ln(A) \sim s$ | 0.90480 | 0.02038 | 0.02327 |
| Exponential, radius, $\ln(A) \sim r$ | 0.91209 | 0.00799 | 0.01819 |
| `nls` Exponential, radius, $A \sim A_0 e^{cr}$ | 0.94799 | <0.00001 | <0.00001 |
| Linear, radius, $A \sim r$ | 0.94862 | <0.00001 | <0.00001 |
| `nls` Exponential, arclength, $A \sim A_0 e^{cs}$ | 0.96113 | <0.00001 | <0.00001 |

| | | | |
|---|---|---|---|
| Quadratic, radius, $A \sim \texttt{poly}(r, 2)$ | 0.96752 | 0.05660 | 0.05660 |
| `nls` Power-law, radius, $A \sim (c_1 r + c_0)^k$ | 0.96780 | 0.05289 | 0.05289 |
| Linear, arclength, $A \sim s$ | 0.97303 | <0.00001 | <0.00001 |
| Quadratic, arclength, $A \sim \texttt{poly}(s, 2)$ | 0.97303 | <0.00001 | <0.00001 |
| Power-law, radius, $\ln(A) \sim \ln(r)$ | 0.97446 | 0.52131 | – |
| Quadratic, $A \sim \texttt{poly}(\theta, 2)$ | 0.97826 | 0.05172 | 0.05172 |
| Exponential, $\ln(A) \sim \theta$ | 0.97936 | 0.49407 | 0.87799 |
| `nls` Exponential, $A \sim A_0 e^{c\theta}$ | 0.98714 | <0.00001 | <0.00001 |
| `nls` Power-law, arclength, $A \sim (c_1 s + c_0)^k$ | 0.98789 | 0.07607 | 0.07607 |

#### S7 Optimal control of planispiral growth

A first step in obtaining the shortest path solution for planispiral growth is to formulate the problem as that of optimal control — defining state, costate, and control variables, boundary conditions, the objective functional, and the Hamiltonian. The state variable is clearly the radius, $r(\theta)$, where winding angle, $\theta$, serves the usual role of a time variable. Boundary conditions for $r(\theta)$ are $(0, r_1)$ and $(\theta_{12}, r_2)$. I will limit the discussion to the case of $r_1$ and $r_2$ separated by a half whorl, $\theta_{12} = \pi$, and $r_2 = r_1 q$, where $q > 1$ is a constant representing the final growth ratio over the entire growth increment.

The quantity to be minimized is arclength over this growth increment $\ell_{12} = \int_{(0,r_1)}^{(\pi,r_2)} r(\theta) \sec \psi(\theta) d\theta$, which defines the objective functional, and establishes the expansion angle, $\psi$, as the control the variable. The optimal control problem seeks a control path, $\psi(\theta)$, that minimizes $\ell_{12}$. The dynamics of the state variable follows

$$\frac{\mathrm{d}r}{\mathrm{d}\theta} = r \tan \psi. \tag{S12}$$

A costate variable, denoted in this section by $p$, has a terminal boundary condition $p(\theta = \pi) = 0$, as there is no terminal reward (i.e., the optimization objective is wholly defined by the objective functional). This leads to the following definition of the Hamiltonian

$$H = r \left( \sec \psi + p \tan \psi \right), \tag{S13}$$

and to the dynamics equation for the costate variable

$$\frac{\mathrm{d}p}{\mathrm{d}\theta} = -\frac{\partial H}{\partial r} = -\sec \psi - p \tan \psi. \tag{S14}$$

Because the Hamiltonian does not depend explicitly on $\theta$ (i.e., is 'time-independent'), its value is conserved along the optimal trajectory. Combined with the terminal condition $p = 0$, I obtain the following results, $H(\theta) = H_0 = \text{const}$ for all $\theta \in [0, \pi]$, $\mathrm{d}p/\mathrm{d}\theta = -c_0/r_2$ at the terminal point, and $H_0/r_2 = \text{const} = \sec \psi(\theta = \pi)$. Rewriting Eq.[S13] as $H = rg(\psi, p)$, I obtain the switching function,

$$\frac{\partial g}{\partial \psi} = \frac{\sin \psi + p}{\cos^2 \psi}, \tag{S15}$$

which determines the slope of the Hamiltonian with respect to the control variable, at given state-, $r$-, and costate-, $p$-, values, and thus the optimal control. Singular control is obtained when the switching function vanishes, $\partial g/\partial\psi = 0$, along a nonzero length segment of $r(\theta)$. Because optimal control seeks to minimize the Hamiltonian, it is clear why singular control and the switching function are of interest.

From Eq.[S15], singular control is obtained when $p(\theta) = -\sin\psi(\theta)$. Substituting into the Hamiltonian (Eq.[S13]), I get $r = H_0 \sec\psi$, and further substituting into state-variable dynamics, Eq.[S12], I finally obtain that singular control, also called *singular arc*, translates to a linearly changing expansion angle,

$$\hat{\psi}(\theta) = \theta - \hat{\theta}, \tag{S16}$$

where $\hat{\theta}$ is a constant parameter. It is not difficult to see that this singular arc defines a straight line, as expected for the shortest path between two points in euclidean space.

However, if the values that the control variable can take are limited, specifically if $\psi \in [0, M]$, a straight line cannot be obtained for the entire optimal path between $(0, r_1)$ and $(\pi, r_2)$. This is a constrained optimal control problem. The optimal path will combine segments of singular control (straight lines), and segments of boundary control; in this case, circular paths ($\psi = 0$) and/or logarithmic spirals ($\psi = M$).

If $q = 1$, i.e. no net growth, $r_2 = r_1$, it is clear that the only possible path is $\psi(\theta) = 0$; a semicircle from $(0, r_1)$ to $(\pi, r_1)$. In all other cases, $q > 1$, either a line or a logspiral segment must be added to the optimal path, to account for nonzero net growth. We already know that, at $\theta = \pi$, $p = 0$ and $\mathrm{d}p/\mathrm{d}\theta = -c_0/r_2 = -\sec\psi_2$ ($\psi_2 = \psi(\pi)$, final expansion angle). We know that on a straight line Eq.[S16] holds, i.e., expansion angle increases linearly. Because $\psi$ cannot go below 0 or above $M$, the winding angle span of a singular-control line segment cannot exceed $M$. Consequently, if $1 < q \leq \sec M$, $H_0 = r_1$, $r_2 = r_1 \sec\psi_2$, $\hat{\theta} = \pi - \psi_2$, and $\psi_2 = \operatorname{arcsec}(q) = \arccos(1/q)$. In other words, the shortest (optimal) trajectory is a circular path of radius $r_1$, from 0 to $\pi - \operatorname{arcsec} q$, followed by a straight line with $r(\theta) = r_1 \sec\psi(\theta)$.

When $q > \sec M$, a terminal line segment, of winding angle span $M$, does not provide the necessary net growth, and an additional terminal logspiral segment is added. In such cases $\psi_2 = M$. On this terminal logspiral, costate dynamics (Eq.[S14]) follows $\mathrm{d}p/\mathrm{d}\theta = -\sec M - p\tan M$, a simple first-order linear ODE with constant coefficients, solved by $p(\theta) = \csc M \left(e^{(\pi-\theta)\tan M} - 1\right)$, after applying the boundary condition $p(\pi) = 0$. As $\theta$ decreases from its terminal values of $\pi$, the costate variable becomes negative and decreases monotonically, until $p(\theta) = -\sin M = -\sin\psi$, the transition point with singular control (switching function, Eq.[S15], vanishes). This is the end-point of a singular arc, a line segment that connects $\psi(\hat{\theta}) = 0$ with $\psi(\hat{\theta} + M) = M$. As before, $r(\hat{\theta}) = r_1$, $r(\hat{\theta} + M) = r_1 \sec M$, and the length of the singular-control line segment is $r_1 \tan M$. Final radius of the line segment, $r_1 \sec M$, serves as the initial condition for the terminal logspiral. Consequently, $q = e^{(\pi-\hat{\theta}-M)\tan M} \sec M$, or $\hat{\theta} = \pi - M - \ln(q\cos M)/\tan M$.

When $\hat{\theta} = 0$, there is no longer an initial circular path included in the optimal solution. That occurs for $q = e^{(\pi - M)\tan M} \sec M$. The optimal solution now begins with a line segment. Finally, for larger growth ratios, $e^{(\pi - M)\tan M} \sec M < q \leq e^{\pi \tan M}$, the initial expansion angle, $\psi_1$, increases as $q$ increases, obeying $e^{\psi_1 \tan M} \cos \psi_1 = q e^{-(\pi - M)\tan M} \cos M$, until the optimal solution is entirely logspiral, $\psi_1 = M$, at the highest possible $q$-value.

The resulting arclength, $\ell_{12}$, of the above optimal-control solutions is given by

$$\ell_{12} = \begin{cases} \pi r_1 & \text{if } q = 1 \\ \left(\pi - \operatorname{arcsec} q + \sqrt{q^2 - 1}\right) r_1 & \text{if } 1 < q \leq \sec M \\ \left(\pi - M - \left(\ln\left(q \cos M\right) + 1\right) \cot M + q \csc M\right) r_1 & \text{if } \sec M < q \leq e^{(\pi - M)\tan M} \sec M \\ \left(-\sin \psi_1 + q \csc M - \cos \psi_1 \cot M\right) r_1 & \text{if } e^{(\pi - M)\tan M} \sec M < q < e^{\pi \tan M} \\ (q - 1) r_1 \csc M & \text{if } q = e^{\pi \tan M} \end{cases} . \tag{S17}$$

Further adding the constraint on diameter of planispiral increment, $r_1 + r_2 = d$ where $d = \text{const} > 0$, provides the initial condition $r_1 = d/(1 + q)$, which can be substituted into Eq.[S17], to provide the ratio of the optimal arclength to that of the unconstrained problem (straight line along the diameter) or to that of a semicircle (i.e., $\psi = 0$), $\ell_{12}/d$ or $2\ell_{12}/\pi d$ respectively. The latter is a convex function of $q \in [1, e^{\pi \tan(M)}]$, with a minimum at $q = 1/\mathrm{W}_0\left(e^{(M - \pi)\tan M + \sec M} \cos M\right)$, $\mathrm{W}_0$ being the zeroth branch of the Lambert W function. This value can be substituted back into Eq.[S17], to obtain the minimum arclength over all values of $q$.

#### S8 Web application

For this paper, I have also written a `WebGL` application to help with creating images of shells, as in Fig. 1*A*,*B*. A snapshot of the code, coinciding with the publication of this report, is available at `https://doi.org/10.5281/zenodo.22165306`.

#### S9 Quote from Blake 1878

Concluding his detailed mathematical analysis of "curves formed by cephalopods and other mollusks", Blake (1878) makes the following observation: "One important value of these equations is to serve as a check on the separate measurements, and thus to gain observations for drawing an average, as there is no doubt that spiral shells are not absolutely geometrically constant, and all measures are therefore more or less approximate."
